# Behavior and Physiology Outpace Form When Linking Traits to Ecological Responses Within Populations: A Meta-Analysis

**DOI:** 10.1101/2025.01.26.634944

**Authors:** Thibaut Rota, Allan Raffard, Iris Lang, Quentin Petitjean, Lisa Jacquin, Olivier Dézerald, Simon Blanchet, Andrew P. Hendry, Régis Céréghino

## Abstract

Intraspecific variability is fundamental to ecology, yet we still know remarkably little about what governs the strength of the associations between traits expressed by individuals and ecological dynamics. To explore this overlooked aspect of diversity, we asked whether the strength of correlations between traits and a wide spectrum of ecological responses could differ (*i*) between intraspecific levels (among *vs*. within populations), (*ii*) among ecological responses across levels of biological organization (from ecological performance to ecosystem functioning), and (*iii*) among trait types (morphology, physiology, and behavior). We performed a meta-analysis synthesizing over a thousand effect sizes from nearly two hundred studies spanning approximately a hundred animal species across a broad range of traits and ecological responses. The average effect size was |*r*| = 0.26 (95% confidence interval: 0.21 – 0.30). At the individual level, effect sizes were larger for ecological performance (foraging, diet) than for fitness (reproduction), and tended to be larger for community responses (e.g., community composition of surrounding organisms). Physiology and behavior showed larger effect sizes than morphology. Our meta-analysis not only confirms that intraspecific trait variability is central to ecological dynamics, but also highlights physiology and behavior as key traits for unraveling the ecological consequences of individual variability.

## Introduction

The question “What do species do in ecosystems?” was raised three decades ago by J. H. Lawton (1994), and has since been foundational to trait-based ecology. Traits, defined as attributes measurable on individual organisms (Violle et al. 2007), describe how organisms interact with each other and with their environment. Trait-based approaches brought the promise to be complementary to taxonomic approaches in that regard (Enquist et al. 2015). Traits can be expressed in physical dimensions (e.g., mass, length, volume, energy, time) that meaningfully reflect variation in the fit of organisms to their environment (Arnold 1983; Violle et al. 2007). As a result, inferences drawn from traits are often transferable to other ecosystems and life forms (Funk et al. 2017). General patterns include allometric scaling between body size and metabolic rate (Kleiber 1932), the leaf economic spectrum (Wright et al. 2004), and *r-*/*K-* life history strategies (Pianka 1970). Further, traits do not have borders: they are often cheap and easy to measure, fostering inclusion among researchers from different backgrounds, as well as traits can describe functional diversity across taxonomic groups, from individuals to ecosystems (Carmona et al. 2016).

### Including Intraspecific Trait Variation in Trait-Based Ecology

Early research in trait-based ecology led to a rethink of how trait-based approaches could improve community ecology (McGill et al. 2006; Petchey and Gaston 2006). However, an important limitation of these early studies was their focus on species’ mean trait values, essentially assuming that all conspecifics share uniform trait values (Albert 2015). This assumption might be convenient, but it is conspicuously problematic, as intraspecific trait variation (i.e., variation in biological attributes among individuals and populations within a species) is a tenet of concepts related to natural selection (Darwin and Wallace 1858), niches (Elton 1927; Hutchinson 1957), and coexistence (Macarthur and Levins 1967). In fact, intraspecific trait variation can be as large as trait differences between species (Lecerf and Chauvet 2008; Siefert et al. 2015; Rota et al. 2022), which presumably matters for species coexistence (e.g., Hart et al., 2016), predator–prey (Toscano and Griffen 2014), and consumer–resource interactions (Raffard et al. 2017; Rota et al. 2018). Intraspecific trait variability is thus increasingly recognized as an important facet of ecology (Bolnick et al. 2011; Violle et al. 2012), provoking discussions on how it may affect food webs (e.g., behavior; Moran et al. 2017) and community assembly (e.g., metabolic traits; Brandl et al. 2022). Despite the scientific opportunities, trait-based studies including the intraspecific component of diversity remains rare (*ca.* 4%; Green et al. 2022). For individualizable organisms, ecological interactions (competition, predation, etc.) are occurring among conspecific and heterospecific individuals, whose traits are similar and different to varying degrees. Therefore, a trait-based approach at the individual level should improve our understanding of such interactions and their emergent properties (Bolnick et al. 2011), whereas neglecting trait variation among individuals can bias inferences (Wong and Carmona 2021). For instance, failing to consider individual trait variability can distort relationships between species richness and functional diversity (Cianciaruso et al. 2009). On the opposite, acknowledging that species trait spaces (the range of phenotypes) evolve in response to competition regimes contributes to a better understanding of diversity–function relationships through the lens of evolution (Barabás et al. 2022).

### Why Assess the Strength of Relationships Between Intraspecific Trait Variation and Ecological Dynamics?

Important insights have emerged from an increased emphasis on intraspecific variation. At local scales, intraspecific and interspecific trait variation have similarly strong effects on community assembly patterns (e.g., species richness and diversity) and ecosystem functioning (e.g., productivity or nutrient cycling) (Des Roches et al. 2018; Raffard et al. 2019). Thereby, answering how much and when the intraspecific level of variation matters has important implications for conservation issues. Indeed, intraspecific trait distributions are altered by human disturbances (Alberti et al. 2017; Sanderson et al. 2023), including population loss (Ceballos et al. 2017), and the erosion of genetic diversity (Exposito-Alonso et al. 2022). Conservationists can take advantage of this line of research in identifying and monitoring particular facets of intraspecific diversity which have implications for ecological functions (Blanchet et al. 2020) and the contributions of nature to people (Des Roches et al. 2021). Understanding the factors that shape variation in the strength of relationships between intraspecific trait variability and ecological dynamics will help to understand how evolution, by affecting traits, can modulate ecological dynamics on similar timeframes (i.e., eco-evolutionary dynamics; Pelletier et al. 2009; Hendry 2017). So far, this research has focused on the ‘*evo*’ aspect of the framework, by asking how trait types (e.g., life history vs. morphology) typically differ in natural selection (Kingsolver et al. 2001, 2012; Kingsolver and Diamond 2011), heritability (Mousseau and Roff 1987), evolvability (Wheelwright et al. 2014), and rates of change (Kinnison and Hendry 2001). As an example, Mousseau and Roff (1987) found that narrow sense heritabilities were higher for morphological traits than for physiological, behavioral, or life-history traits, whereas Kingsolver et al. (2001) found that selection in natural populations was stronger on morphological traits than on life history or phenological traits. We will instead focus here on the ‘*eco*’ aspect, where particular trait types could imply different strengths of relationships with ecological processes, with an example being the special role that behavior could play on ecological dynamics (e.g., niche construction; Laland et al. 1999; Laland 2004).

To date, studies have examined the ecological impacts of populations that differ in key traits (e.g., gill raker morphology in anadromous or landlocked fish populations; Post et al. 2008), but we are far from a broad-scale picture of how different factors (e.g., trait types, types of ecological responses, ecological contexts, levels of biological organization) can modulate how individual variability in phenotypic traits affects ecological dynamics (Gibert et al. 2015). For instance, the extent to which individual-level trait variability exhibits similar effects on ecological processes as the trait variability observed among populations remains unclear. On the one hand, we could expect population-level trait variability to show stronger relationships with ecological dynamics than individual-level trait variability, since evolutionary processes such as local adaptation are expected to drive population-level divergence in average trait values (Post et al. 2008). On the other hand, trait variation among individuals within populations has shown similar levels to trait variability among populations (e.g., Messier et al. 2010; Rota et al. 2024), perhaps as a result of metapopulation dynamics contributing to gene flow and spatial homogenization of biological attributes.

How the strength of relationships between traits and ecological responses varies across increasing levels of biological organization, from individuals to ecosystems, is still debated (Violle et al. 2007; Enquist et al. 2015; Chacón-Labella et al. 2022). Some authors predict stronger relationships for responses at low levels of biological organization (i.e., the responses proximal to organisms, such as food intake and growth rate) than at high levels of organization (i.e., responses distal to the organisms, or involving more indirect links, such as community structure and ecosystem functioning; Bailey et al. 2009). Others have suggested an opposite pattern, with strong relationships at community and ecosystem levels (Des Roches et al. 2018; Raffard et al. 2019).

### The Ecological Functionality of Traits

In ecology, we generally expect that the traits we study are functional; that is, linking to ecological performance or fitness (Arnold 1983; Violle et al. 2007). Yet the *functionality of traits* remains a matter of debate, as it affects ecological understanding and the choice of traits used by ecologists (Mlambo 2014; Lefcheck et al. 2015; Dawson et al. 2021). In biological terms, all traits are functional because they link to one or several biological functions, constituting functional living entities (Sobral 2021); yet the definition of a functional trait is discipline-specific (e.g., see Dawson et al. 2021 and Sobral 2021). Rather than discussing these definitions, we propose to investigate here what drives *variation in the strength* of relationships between traits and various ecological parameters. For example, a meta-analysis showed relatively small absolute correlation strengths (i.e., absolute correlation coefficients |*r*|) between animal behavior and fitness (Moiron et al. 2020). Even so, behavioral traits can be functional for other reasons. For example, individual variability in activity levels is often strongly associated with energy intake, growth rate (Biro and Stamps 2008), and with the behavioral type of foraged prey (McGhee et al. 2013), which can alter community and ecosystem processes (LaBarge et al. 2024; Szangolies et al. 2025). Understanding whether a trait consistently shows strong associations with diverse ecological responses across many taxa (what we define here as *ecological trait functionality*) can better inform ecologists, improve trait-based inferences at the individual level, and guide future research on this often-overlooked aspect of diversity.

A particularly important functional trait is body size. Body size is used in the majority of trait-based studies (Green et al. 2022). It can span several orders of magnitude across and within taxa, and it explains many processes with great predictive power (Peters 1993). Nevertheless, for animal species exhibiting indeterminate growth, it may not be always meaningful to explore associations between body size and fitness among individuals of different age classes (Thompson and Fincke 2002), since comparing life histories and fitness across ontogenetic stages can be trivial (e.g., between individuals at the immature and reproductive stages). Furthermore, functional diversity is highly multidimensional (de Bello et al. 2021), and it is unlikely that body size alone can explain all the ecological variation embedded among individuals. For example, large differences exist in morphology (Post et al. 2008), physiology (Careau et al. 2014) and behavior (Sih et al. 2004) between similarly sized and aged individuals (Niemelä and Dingemanse 2018*a*). However, we do not know yet if differences exist in the strength of relationships that these types of traits can maintain with fitness or other important ecological responses (i.e., their ecological functionality). It is time to look beyond body size to improve our understanding of the ecological implications of individual trait variation (Gordon 2011; Toscano et al. 2016; Moran et al. 2017; Brandl et al. 2022; Schleuning et al. 2023).

Morphology, physiology, and behavior are different facets of a phenotype. Morphology reflects the abilities of an individual to perform in its environment given biomechanical constraints (Van Valen 1965; Arnold 1983). We can also tie indirect links between morphology and energetics. For instance, the relative mass of metabolically active organs is related to energy expenditures (Careau et al. 2008). Many studies have investigated the morphological variation that occurs between individuals specializing in different sets of resources (e.g., Bolnick et al. 2007), with the idea that variability in the form of key trophic apparatus, such as beak size in Darwin’s finches, conditions the diet of individuals and thus their evolution (Van Valen 1965; Boag and Grant 1981). Physiology expresses more direct energetic currencies, such as metabolic rates (i.e., energy loss), which link closely to ecological performance, including feeding rate (i.e., energy intake) and growth rate (i.e., available energy not allocated to maintenance or reproduction; Biro and Stamps 2008; Careau and Garland 2012). Through trophic and competitive interactions, physiological traits may be involved in community assembly (Brandl et al. 2022). In parallel, a large body of research suggests that behavioral traits are particularly important in ecology (Gordon 2011; Toscano et al. 2016). Individual variation in behaviors such as activity and boldness can strongly affect the outcomes of predator-prey interactions (McGhee et al. 2013). Behaviors such as activity, aggressiveness, or boldness can express syndromes (Sih et al. 2004) involving energetic trade-offs (Careau et al. 2008). Behavioral syndromes can include covariations with metabolic rates and hormone expression, linking them to broader syndromes such as the ‘pace-of-life syndrome’ (POLS), which posits a continuum from slow to fast life histories (Ricklefs and Wikelski 2002; Réale et al. 2010; Wright et al. 2019). Therefore, we expect that physiological and behavioral traits, as well as other energy and matter currencies (e.g., elemental stoichiometry; Elser et al. 2000), exert strong links with ecological performance, fitness, and hence ecological dynamics (Gordon 2011; Moran et al. 2017; Brandl et al. 2022). Despite available qualitative examples on how individual variation in the traits of animals affects various ecological processes, no clear pattern has emerged (fig. 1 and table 1), and what drives the strength of those relationships remains unanswered.

**Figure 1:**
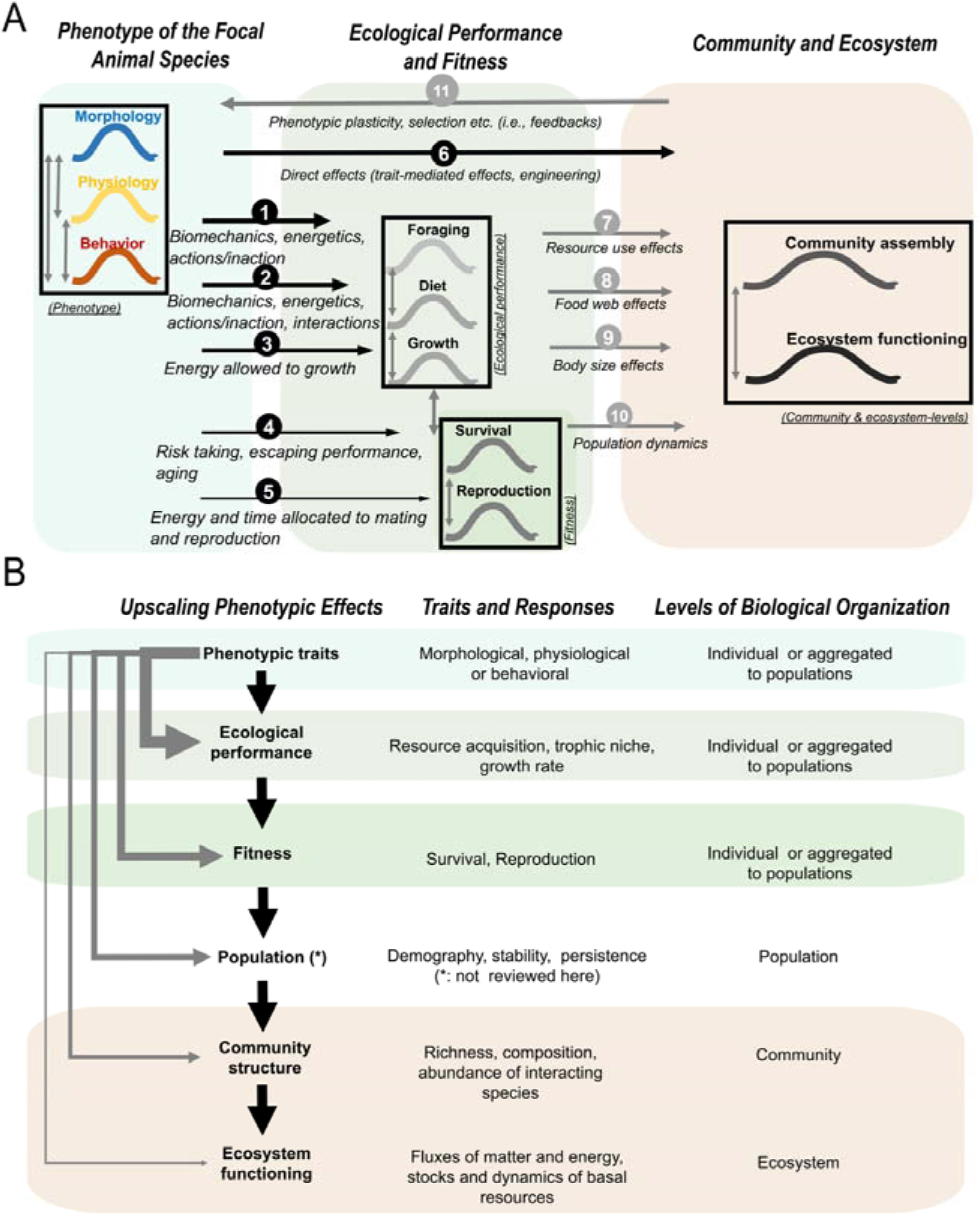
(A) Conceptual diagram showing how phenotypic trait variability (morphology, physiology, behavior) relates to various ecological responses in animals. Black arrows show links that are relevant to this meta-analysis, with real-world examples of relationships depicted in table 1, where numbers in parentheses correspond to each link of black arrows shown in fig. 1A. With black arrows shrinking and elongating, we represented how complexities increase when a phenotypic trait links from proximal to distal responses, i.e., how much an individual eats, to what it eats, to how much it grows, to its odds of survival, and if alive, to its success in reproduction. In that sense, we think of the mechanisms underlying *links #1* to *#5* as cumulating their complexity (i.e., *link #5* involves more complexity than *link #1*). *Link #1* may be the most straightforward (feeding rate could be explained by energetic needs, search rate, and handling time). Fitness responses, however, in addition to integrating the complexities of *links #1–4*, add those related to energy and time investment in mating and reproduction. *Link #6* shows direct links from the phenotype to community and ecosystem functioning (e.g., trait-mediated effects). *Links #7* to *#10* show how effects on ecological performance and fitness (e.g., differences in terms of per capita utilization of resources, diet, etc.) can indirectly affect communities of surrounding species and ecosystem processes (fluxes of energy and matter). Change in ecological conditions can induce phenotypic plasticity or microevolution, triggering feedbacks on the phenotype (*link #11*). However, the *links #7–10* and *#11* (gray arrows) have not been synthesized quantitatively here. Double vertical arrows depict covariations. (B) The strength of relationships between phenotypic traits and ecological responses is expected to change across levels of biological organization. Black arrows depict intermediate relationships propagating phenotypic effects across biological levels of organization. Following hypothesis 2 (see main text), the strength of relationships between phenotypic trait variation and ecological responses could decrease (width of gray arrows) with increasing levels of biological organization (Bailey et al. 2009), from responses located on the focal animal itself (top), to community and ecosystem levels (bottom). A likely alternative is that phenotypic traits affect directly the communities of organisms (*link #6* in fig. 1A). All relationships depicted by gray arrows in fig. 1B are evaluated in this meta-analysis, except the one for population (*).

**Table 1:** Chosen examples of traits-to-ecological response relationships from studies constituting our dataset. Studies include measures made on individuals (‘ind’), populations (‘pop’), and colonies of social insects (‘col’). The numbers in bold and in brackets correspond to *links* shown in fig. 1A.

| Level | Trait(s) | Trait type | Species | Ecological response(s) | Trait-to-function relationship (hypothesized or observed) | References |
| --- | --- | --- | --- | --- | --- | --- |
| ind | Tail shape | Morphology | *Hyla chrysoscelis* (amphibian) | Survival when exposed to predators (fitness) | Induced changes in the tail shape of tadpoles exposed to predators (**11**) enhanced their survival, affecting their escaping abilities (**4**) | McCollum and Van Buskirk 1996 |
| ind | Mandible shape | Morphology | *Gammarus fossarum* (arthropod) | Litter consumption rate (foraging performance) | The shape of mandibles did not alter the performance of the amphipods to consume leaf litter (**1**) | Rota et al. 2018 |
| ind | Functional morphology | Morphology | *Micropterus salmoides* (fish) | $\Delta C^{12,13}$ and $\Delta N^{14,15}$ (trophic niche) | Beyond large effects of ontogeny, no relationship between morphological traits and trophic niches was detected (**2**) | Zhao et al. 2014 |
| ind | Benthic vs. limnetic morphotypes | Morphology | *Gasterosteus aculeatus* (fish) | Invertebrate community structure (community) | Benthic and limnetic morphotypes are specialized in different prey, inducing effects on community structure (**2, 8**) | Des Roches et al. 2013 |
| pop | Gill raker morphology (anadromous vs. landlocked ecotypes) | Morphology | *Alosa pseudoharengus* (fish) | Chlorophyll a (ecosystem functioning) | Anadromous and landlocked ecotypes feed on different zooplankton species according to their gill rakers morphology, thereby altering biomass of phytoplankton (**2,8**) | Post et al. 2008 |
| ind | Corticosterone levels | Physiology | *Thalassarche melanophrys* (bird) | Number of chicks (fitness) | Elevated corticosterone levels (stress hormone) were associated with lower reproductive output (**5**) | Angelier et al. 2010 |
| ind | Mass-specific standard metabolic rate | Physiology | *Desmognathus brimleyorum* (amphibian) | Feeding rate on flies (foraging performance) | Individuals with a higher mass-corrected metabolic rate consumed prey at faster rates (**1**) | Gifford et al. 2014 |
| ind | Elemental stoichiometry (C, N, P) | Physiology | *Procambarus clarkii* & *Faxonius limosus* (arthropoda) | $\delta^{13}C$ and $\delta^{15}N$ (trophic niche) | Elemental imbalances (C, N, P) between consumers and resources drive the diet of consumers (**2**) | Lang et al. 2021 |
| ind | Mass-specific standard metabolic rate | Physiology | *Phoxinus phoxinus* (fish) | Pelagic production (community & ecosystem functioning) | Higher energy expenditures altered the community structure (**1,2,6,7**) and the pelagic production of phytoplankton | Raffard et al. 2021 |
| col | Boldness, activity | Behavior | *Apis mellifera* (arthropoda) | Colony weight, survival after winter (fitness) | Colonies that expressed active and bold behavior accumulated more reserves and grew faster, increasing winter survival (**1,3,4**) | Wray et al. 2011 |
| ind | Social dominance | Behavior | *Lepidodactylus lugubris* (squamate) | Prey feeding rates (foraging performance) | Socially dominant geckos consumed more crickets than subordinate individuals (**1**) | Short and Petren 2008 |
| ind | Activity | Behavior | *Esox lucius* (fish) | $\delta^{15}N$ (trophic position) | Active pikes had higher trophic positions (**2**) | Nyqvist et al. 2018 |
| ind | Activity | Behavior | *Phoxinus phoxinus* (fish) | Algal biomass and litter decomposition rate (community & ecosystem functioning) | Active minnows consumed potentially more grazers but fewer detritivores (**1,2**), increasing algal production and decomposition rate (**6,7,8**) | Raffard et al. 2023 |

### Questions and Hypotheses

We addressed the above topics through a meta-analysis of a large diversity of traits, systems, and species in animals (fig. 1A and table 1). Specifically, we explored the strength of correlations between intraspecific trait values and a large array of ecological responses (fig. 1B). To go beyond the already well understood influence of body size, we compared individuals that differed little in size, or for which the potential dependence of traits on body size or ontogeny has been accounted for statistically. After first testing if the overall strength of relationships departs from zero (*H0*), we asked three additional questions.

(*Q1/H1*) Does the strength of relationships between traits and ecological responses (hereafter ‘effect sizes’) differ depending on whether they were assessed within populations (***i_a_***) versus among populations (***i_b_***)? We expected that ***i_a_* < *i_b_***, because individuals should show stronger phenotypic divergence among- than within populations. An alternative expectation was that ***i_a_* ∼ *i_b_***, if, for instance, the populations sampled to obtain individual-level estimates (***i_a_***) had large sizes or were a part of interconnected populations, making them phenotypically representative of the entire metapopulation (***i_b_***) (Messier et al. 2010; Rota et al. 2024).

(*Q2/H2*) How do effect sizes vary among ecological responses at different scales of biological organization? We ranked ecological responses from a lower biological scale (***ii_a_***), i.e., proximal responses to the focal individuals (e.g., foraging performance, growth, diet, and fitness), to a higher biological scale (***ii_b_***), i.e., distal responses from the focal individuals including community assembly (e.g., community structure, diversity, biomass of heterospecifics; further abbreviated ‘community’) and ecosystem functioning (e.g., nutrient cycling, rates of energy and matter flow of basal resources; further abbreviated ‘ecosystem’; see fig. 1B). We expected that effect sizes would decrease from proximal to distal responses (***ii_a_* > *ii_b_***; as the width of gray arrows in fig. 1B). Our rationale was that the strength of relationships between traits and responses should dissipate as the number of intermediate relationships increases (i.e., with the sum of black arrows in fig. 1B; see the algebraic rationale in Bailey et al. 2009). On the contrary, effect sizes could be stronger for distal than for proximal ecological responses (***ii_a_* < *ii_b_***) if the strength of intermediate relationships shows additivity or multiplicativity through biological scales (Bailey et al. 2009), or if direct, trait-mediated interactions prevail (*link #6* in fig. 1A; Schmitz et al. 2004).

(*Q3/H3*) How do effect sizes vary among trait types? Effect sizes were divided into (***iii_m_***) morphology (e.g., body shape, relative length of functional body parts), (***iii_p_***) physiology (e.g., metabolic rate, hormone levels, elemental stoichiometry), and (***iii_b_***) behavior (e.g., activity, boldness, sociability). We hypothesized that behavioral and physiological traits would show stronger correlations with ecological responses than would morphology (i.e., ***iii_m_ < iii_p_ ∼ iii_b_***). Behavior expresses what an individual *does*, and so we expected that behavioral traits should be good proxies for interactions among organisms, and thus for their ecological differences. Physiology expresses internal biological rates that are often linked to energetics, behavior, and life history strategies, and should thus also show strong ecological effects. Morphology, by contrast, poses physical constraints on what an individual *can* do, and so it has been argued that behavior and physiology would be closer to the ecologies of animals than would morphology (Wainwright and Reilly 1994). This idea is also at the core of organic selection theories (Baldwin 1897), which often suggest that behavior, through nongenetic inheritance, Baldwin effects, and/or niche construction, is a main driver of the evolution of the organisms and other characters (physiology and morphology), and of the eco-evolutionary feedbacks of those changes (Laland et al. 1999; Hall 2001; Danchin et al. 2011). We expected, however, that morphological differences might be just as important as other trait types when individuals have been diverging from hundreds to thousands of generations and for which ecological differences are strong (e.g., ecological radiation between limnetic and benthic ecomorphs within a single lake population; Harmon et al. 2009).

We tested *H2* across all trait types, and similarly, we tested *H3* across all response types, and both hypotheses were tested at the individual level, so that we benefited from the maximal sample size (effect size estimates) to test each hypothesis while accounting for the multi-level nature of the data (see Methods). To facilitate comparison to our global estimate of the strength of correlations between individual trait variation and ecological responses beyond body size, we compiled an ad hoc dataset on the strength of correlations between body size varying among individuals and various ecological responses (Supplementary Materials, Section 2). We expected that the effect sizes for body size would be much larger than those for morphological, physiological, and behavioral traits that varied independently of body size or ontogeny (or whose dependence on size was accounted for), given the strong scalings that body size maintains with energetics, stoichiometry, and predator-prey interactions (Brown et al. 2004; Brose et al. 2006; Vanni and McIntyre 2016), to cite only a few ecological domains where body size matters (Peters 1993).

## Methods

### Search Strings and Selection Criteria

We searched for 30 keyword combinations in Web of Science (‘all databases’), Scopus, and Google Scholar (Supplementary Materials, Section 1; table S1), through which we obtained a total of 6904 studies. After having manually sorted duplicates in Excel (n = 335), we obtained 6569 studies, of which 613 followed our scope after having read titles and abstracts. Our scope was studies at the intraspecific level, in animals, that investigated relationships between individual differences in traits and various ecological responses. Exclusion criteria at this stage were studies focusing on groups other than animals, investigating questions other than traits-to-ecological responses, or studies conducted at the interspecific level.

After a full-text examination of each 613 studies, 121 met our criteria. We considered empirical studies (*i*) on wild animals, or wild animals that had been reared in the laboratory (in rare cases under captive conditions for a few generations), with observations conducted either in the field, microcosms, or mesocosms. (*ii*) Studies reporting correlation coefficients, or other statistics, for relationships between a phenotypic trait measured among several individuals (or averaged among populations or colonies of eusocial animals) and a measured ecological response. The trait types that we considered were (*iii*) morphological, physiological, or behavioral, regardless of whether the traits were categorical or continuous. (i*v*) We did not include ecological responses reflecting population dynamics or population persistence, because these are long-term phenomena for which relationships with trait variability are often difficult to estimate empirically. (*v*) We were interested in studies focusing on the identity of phenotypes (i.e., one phenotype versus another), not diversity (i.e., statistics from treatments that included mixtures of phenotypes were excluded). (*vi*) We did not define life history as a trait type, since we categorized its components as ecological responses (e.g., growth, survival, and reproduction). Finally, (*vii*) we focused on traits beyond body size (see our motivation in the Introduction) by selecting studies that considered similarly sized, and/or aged individuals, or that accounted for the potential influence of body size or ontogeny on correlations (for a detailed description of choices made by authors regarding body size and ontogeny, see Box S1 in Supplementary Materials, and publicly available data). From the choices made by authors, we gave a score for each study representing our confidence that the relationship was independent from body size variation. This confidence score regarding independence from body size (high score) or potential bias regarding the presumed lack of independence from body size and reported statistics (low scores) was tested as a moderator.

We finally scrutinized the references of these 121 papers that we retained, and added 49 additional studies meeting our selection criteria. We further added 21 studies from our personal libraries (fig. S1). Studies reporting undetailed statistics (e.g., only *P*-values) or those recently retracted were excluded from the final dataset. However, we kept the effect sizes (*r* values) from Bolnick and Paull (2009) even though they retracted their paper, since the error was an incorrect calculation of the *P-*values, which did not affect the validity of the *r* coefficients, hypotheses, or methods.

### Intraspecific Level, Trait Types, Ecological Responses, and Covariates

Related to our first hypothesis (*H1*), we noted the intraspecific level from which the relationships belonged (i.e., individuals, populations, or colonies of eusocial animals). Please note that while we may refer to the among-individual level, most effect sizes at the individual level were actually encapsulating intra- and inter-individual components of trait variation, and therefore are better understood as whole *phenotypic correlations* with ecological responses (Niemelä and Dingemanse 2018*b*).

Related to our second hypothesis (*H2*), we considered the following seven types of ecological response: ‘*foraging efficiency*’ (resource consumption or feeding rates); ‘*trophic niche*’ (trophic niche position, trophic level, and degree of specialization inferred from stable isotopes or diet); ‘*growth*’ (size-standardized growth rates over time based on body length or body mass); ‘*survival’* (survival rates obtained from capture-mark-release-recapture in the wild, or from controlled experiments); *‘reproduction’* (mating, egg numbers, clutch sizes, or sibling survival); ‘*community*’ (abundances, biomasses, diversity, and community composition of organisms of at least two species interacting with the focal phenotypes); and ‘*ecosystem*’ (standing stocks or dynamics of basal resources such as primary producers or detritus, as well as energy and matter cycling such as C and N cycling).

In relation to our third hypothesis (*H3*), we categorized traits into morphology, physiology, and behavior. However, original traits were more diverse than those three categories. Morphological traits included body shape, trophic apparatus shape, linear morphometrics of specific features, locomotion traits, coloration, body condition, relative size of reproductive organs, relative size of brain, and morphotypes. Physiological traits were assimilation, energy reserves, excretion, metabolic traits, elemental stoichiometry, hormone levels, immunity, or physiological syndromes. Behavioral traits, following their methodological description in primary studies, were assigned to one of the five axes of animal personality: aggressiveness, exploration, boldness, sociability, and activity (see definitions and methodological examples in table 3 of Réale et al. 2007), to which we added another category of traits reflecting cognitive abilities of animals (e.g., problem-solving). We decided between activity and exploration using the definitions of Réale et al. (2007). All behavioral traits fell naturally within these categories. As moderator covariates (that we tested individually on separate models), we considered the trophic level (i.e., primary consumers feeding on basal resources vs. predators feeding on other animals), and the ecosystem type (i.e., aquatic vs. terrestrial). In assigning species with aquatic-terrestrial life cycles to either one of those realms, we relied on the stages investigated by the authors, e.g., aquatic for tadpoles and terrestrial for frogs. Finally, we considered the methodological approach (i.e., field observations, mesocosm, or microcosms), and the year of publication.

### Calculation of Effect Sizes

We collected correlation coefficients (*r*) or other statistics (e.g., *F* or *t* statistics) corresponding to each trait-to-ecological response relationship. Original estimates were transformed to correlation coefficients *r* and then to Fisher’s correlation coefficients *Zr,* following established procedures (Nakagawa and Cuthill 2007; see table S2). Since we aimed to quantify the strength of correlations between intraspecific trait values and ecological responses, the direction of relationships was not meaningful. We therefore used absolute correlation coefficients |*Zr*| in our analyses – see also Bailey et al. (2009). For each effect size |*Zr*|, we computed a sampling variance (*vi*) that we used in our models detailed below (Nakagawa and Cuthill 2007; Nakagawa et al. 2022).

### Statistical Analyses

We performed statistical analyses using R version 4.3.2. (R Core Team 2023). We used hierarchical multi-level models that account for phylogeny (function ‘*rma.mv()*’ in the R package ‘*metafor*’; Viechtbauer 2010, see the package’s website: https://www.metafor-project.org/doku.php/metafor). These models take random effects accounting for between-and within-study variance components, as well as a matrix of phylogenetic relatedness among species in the error structure. Our models addressed the three usual sources of nonindependence of meta-analyses in ecology and evolution (Noble et al. 2017; see also a guide by S. Nakagawa and M. Lagisz (2016) here: https://environmentalcomputing.net/statistics/meta-analysis/meta-analysis-3/). In addition, we modeled a variance – covariance matrix *V* reflecting the nonindependent structure of sampling variances (noted “*vi”* hereafter) in our dataset, and obtained cluster-robust estimates regarding selective reporting bias (see Yang et al. 2024*a*).

We computed phylogenetic relatedness among species following Moran et al. (2021). We extracted phylogenetic and taxonomic information from the Open Tree of Life (https://tree.opentreeoflife.org/; Hinchliff et al. 2015). We then resolved any polytomies by randomization in the R package ‘*rotl*’ (Michonneau et al. 2016). Branch lengths were estimated using Grafen’s method (Grafen 1989) in the R package ‘*ape*’ (Paradis and Schliep 2019), and we added a residual error term that accounts for the computed phylogenetic correlation matrix in our meta-analytic models. We accounted for within-study nonindependence by adding a within-study random effect term (nested within *studyID*), attributing an identifier to all effect sizes (see Noble et al. 2017; Yang et al. 2024*a*). Selective reporting bias and other sources of nonindependence inherent to the structure of the data were accounted for by modeling an appropriate sampling variance – covariance matrix *V*, and by following a two-step cluster-robust estimation procedure (Yang et al. 2024). We computed *V* with the function *vcalc()* of the package ‘*metafor*’, accounting for the basal level of our observations (*effect sizeID*), the cluster of the model (*studyID*), and a subgrouping variable with respect to trait-to-response combinations (n= 468) (subgroup argument), setting covariation among subgroups as ρ=0.6. This accounted for the fact that effect sizes share means and errors within those combinations, while acknowledging the multi-level structure of the data. For each model (see fixed-effect structures below), we re-ran the output from the function *rma.mv()* to get a cluster-robust estimation of effect size estimates and their variance using the function *robust()* of the package ‘*metafor*’ (i.e., two-step procedure described in fig. 7 in Yang et al. 2024*a*).

Sampling variance *vi* and mean-centered year of publication are indicators of small study effects and publication time-lag bias, respectively (Nakagawa et al. 2022). We evaluated and accounted for these two sources of bias by adding these terms as covariates in our models, so effect size estimates are given for *vi* = 0, and for the average year in our dataset which was 2012.6 (Nakagawa et al. 2022). Using the same error and random-effects model structure as mentioned above, we first estimated the grand-mean effect size with an intercept-only model, to test if effect sizes depart from zero (*H0*). We then set the intraspecific level as a fixed effect (*H1*). Here we tested if effect sizes at the individual-level were lower than those at the population-level, where we merged ‘populations’ and ‘colonies of eusocial animals’ to reduce the imbalance of effect sizes at the individual (n=952) and population levels (n=59). We then tested our hypotheses regarding differences among trait types and ecological response types (*H2* and *H3*) on the individual-level dataset, with n=952 effect sizes (94% of the dataset), since our main focus was on the individual level. We also tested *H2* and *H3* using the entire dataset to see if adding estimates at the population level would change our conclusions formed at the individual level (see table S3). For the second hypothesis (*H2*), we used a model including ecological response as a fixed effect and trait type as a random effect. We then tested *H3* by adding trait type as a fixed effect, and ecological response as a random effect. For all hypotheses, we considered statistical significance among groups with comparison tests using the *glht()* function of the package ‘*multcomp*’ (Hothorn et al. 2008) accounting for the multi-level structure of the dataset, with an alpha threshold of 5%. For *H2,* we had no main a priori assumption about the direction of differences among ecological responses (we hypothesized either a decrease or an increase across increasing levels of biological organization), and so we performed two-tailed Tukey multi-comparisons for *H2*.

Our hypothesis for *H1* was that the effect sizes at the population level would be larger than those assessed at the individual level. For *H3*, we hypothesized that both physiological and behavioral traits would exert stronger correlations with ecological responses than would morphological traits. Therefore, for *H1* and *H3,* we tested one-tailed planned contrasts [populations > individual-level for *H1*, physiology/behavior > morphology for *H3*, respectively, using *alternative= “greater”,* in *glht()*]. We also tested two-tailed planned contrasts for *H1* and *H3*. We computed pseudo-*R²* values as the proportional reduction of the sum of the variance components between a null model without the fixed effect of interest (intraspecific level for *H1*, response type for *H2,* and trait type for *H3*) and a model with the relevant fixed effect, on models refitted using maximum likelihood estimation (Viechtbauer 2010). We then tested the whole significance of each of these three fixed effects by comparing each pair of model (with or without the fixed effect), using a likelihood ratio test (LRT; Viechtbauer 2010).

Converting correlation coefficients *Zr* to their absolute values |*Zr*| can cause |*Zr*| to be biased upward, since when *Zr* is close to zero, any estimation error will increase *|Zr|* (Morrissey 2016). Furthermore, since |*Zr*| ≥ 0, we cannot directly test its departure from zero (*H0*). To correct our effect size estimates for these two issues, we generated null effect sizes |*Zr|*_null_ (Raffard et al. 2019). For each effect size in our dataset, we randomly simulated a distribution of 1000 *t* statistics based on sample sizes *N* of each effect size under the null hypothesis of no correlation. We then converted these *t* statistics into |*Zr*| values and averaged them for each effect size. Using the same model structure as for each of the models we used (from models *H0* to *H3*), we estimated |*Zr|*_null_ effect sizes. We obtained unbiased |*Zr*| estimates and their 95% confidence intervals by subtracting the estimated |*Zr|*_null_ from the modeled estimates of |*Zr|*. We back-transformed |*Zr|*_unbiased_ to |*r*|_unbiased_ effect sizes to report a popular statistic of association strength in ecology.

As explained above, we accounted for selective reporting bias with a cluster-robust estimation. We also assessed publication bias with a funnel plot on the residuals of |*Zr|* from the intercept-only model including the whole dataset (*H0*). We computed total heterogeneity and tested its significance with a *Q*-test on this same model, as we computed heterogeneity between- and within-study, and phylogenetic heterogeneity (*I*²) using Viechtbauer’s method (Viechtbauer 2010; see R code associated with the paper).

We visualized raw effect size distributions and model estimates with orchard plots (package ‘*orchaRd*’; Nakagawa et al. 2023), but modified the function so that the lower prediction and confidence intervals at 95% were bounded to zero, as by definition, absolute effect sizes cannot extend below zero. Our alignment with the PRISMA guidelines for reporting meta-analysis in ecology and evolution (O’Dea et al. 2021) is shown in Supplementary Materials, Section 1, table S4.

## Results

Our dataset comprised 1011 effect sizes from 187 studies, covering 126 animal species across seven major taxonomic groups (fig. 2A), and a wide range of ecological contexts (fig. 2B). Morphology, physiology, and behavior were represented by 352, 205, and 454 effect sizes, respectively. Effect sizes expressed ecological relationships at the individual level (94%), population level (4%), or among colonies of eusocial animals (2%).

**Figure 2:**
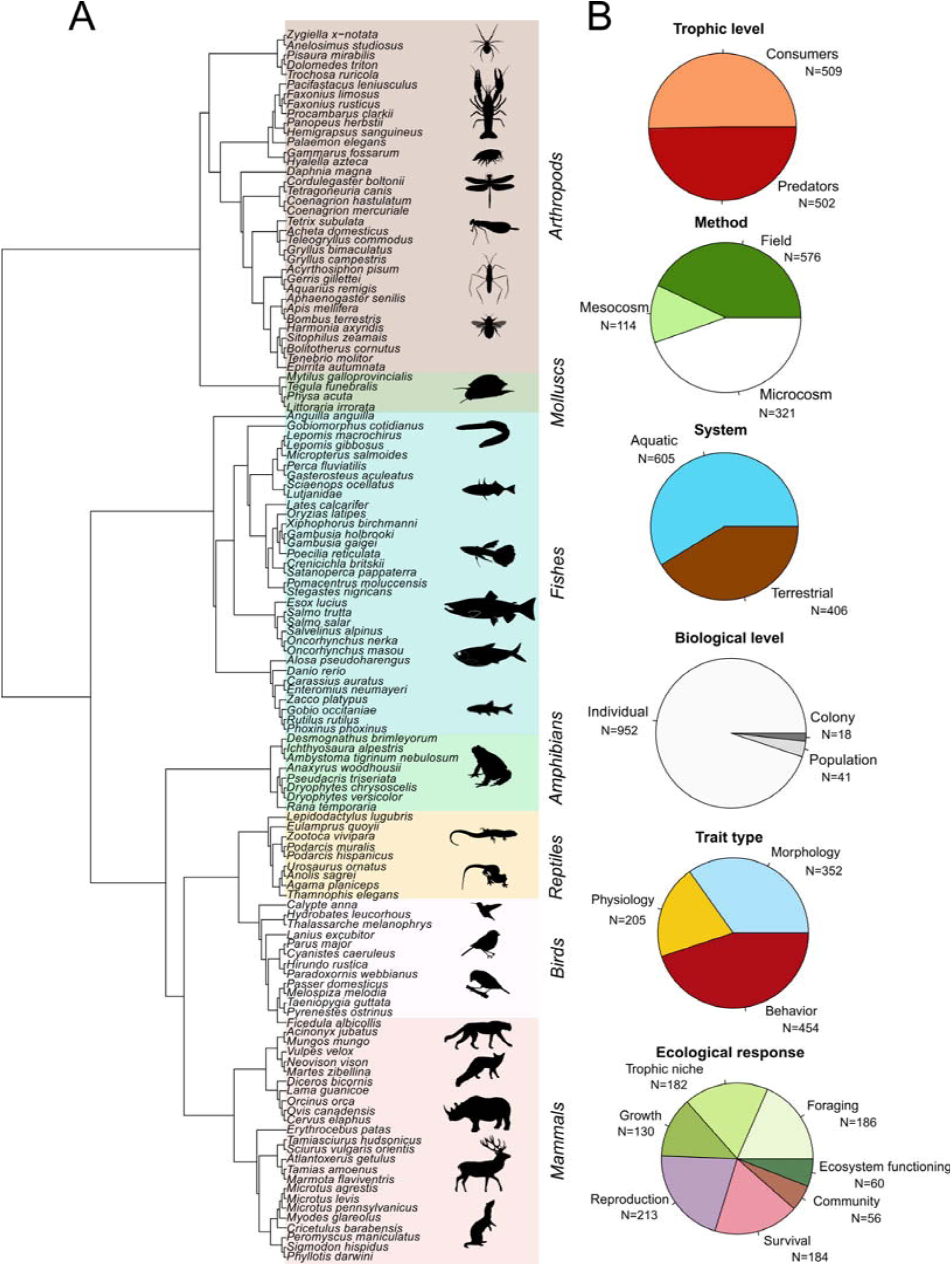
(A) Phylogenetic tree of species in our dataset, with species names and taxonomic groups and silhouettes of some taxa within each group. (B) Pie charts of main factors in our meta-analysis, with respective sample sizes of |*Zr|* effect sizes.

### Heterogeneity and Covariates

The total heterogeneity in our intercept-only model was high (88%; *Q_df=_*_1008_ = 6256.7; *P* < .0001). Phylogeny accounted for a surprisingly large amount of the total heterogeneity in effect sizes (15%). Most of the heterogeneity was within studies (44%) rather than among studies (28%). We did not observe any effect of our covariates, and accordingly, distributions of effect sizes were indistinguishable between systems (aquatic *vs.* terrestrial; *z* = 0.84; *P* = .402), trophic levels (consumers *vs.* predators; *z* = 1.04; *P* = .298), and methods (field observations, mesocosms, and microcosms; *z* < 1.01; *P* > .312; Supplementary Materials, fig. S2–S4). It is noteworthy that the two levels – ‘colonies of social animals’ and ‘populations’ – that we merged to test for differences between intraspecific levels (among-population vs. within-populations) shared overlapping effect sizes on average (fig. S5).

### Publication Bias

The funnel plot of residuals from the intercept-only model was roughly symmetric, with the majority of points falling within the pseudo-confidence region at 95% (fig. S6), suggesting only minor publication bias. We did not observe temporal bias (*t* = –1.64; *P* = .1021); but, as we worked with absolute effect sizes, the effect of the sampling variance (*vi*) was strongly positive (*t* = 5.25; *P* < .0001). We did not find any association between the body size independence confidence score assigned to estimate the degree of independence from body size (high scores) or, conversely, the potential influence of body size (low scores) on the |*Zr*| effect sizes reported here (*t* = 1.06; *P* = .291; fig. S7).

### Estimated Effect Sizes and Results Regarding Hypotheses

The global effect size (*H0*) depicting the strength of relationships between traits varying among individuals or populations, and ecological responses, was different from zero (|*r*|_unbiased_ = 0.26), as its 95% confidence intervals (further abbreviated ‘95% CI’) excluded zero (0.21 – 0.30; table 2).

**Table 2:** Mean effect sizes and 95% confidence intervals for (A) the intercept-only model (global estimate, *H_0_*), and the model for intraspecific levels (among- and within populations; *H_1_*) for the entire dataset. (B) Estimates given for response types (*H_2_*) and trait types (H3) are based on a subset of the data, including only effect sizes at the individual level (n=952; 94% of the dataset). Estimated effect sizes are given as |*Zr*| and |*r*|, as well as with their unbiased estimates (|*Zr*| – |*Zr*| _null_ and |*r*| – |*r*| _null_). Confidence intervals at 95% are given in brackets. For each model, (**†**) indicates the category with the lowest estimated effect size, and the categories shown in bold (**\***) are those with significantly higher estimates compared to (**†**). Pairwise comparison statistics (*z*- and *P*- values) are given accordingly (two-tailed tests for response type differences in *H2*, one-tailed tests for intraspecific level differences in *H1* and for trait type differences in *H3*).

| Parameters (/model) | $\vert Zr\vert $ | $\vert r\vert $ | $\vert Zr\vert _{\text{null}}$ | $\vert Zr\vert _{\text{unbiased}}$ | $\vert r\vert _{\text{unbiased}}$ | $z$ | $P$ |
| --- | --- | --- | --- | --- | --- | --- | --- |
| **(A) (/Intercept-only)** |  |  |  |  |  |  |  |
| Global estimate | 0.32 (0.26 – 0.37) | 0.31 (0.26 – 0.35) | 0.05 (0.05 – 0.06) | 0.26 (0.21 – 0.31) | 0.26 (0.21 – 0.30) | – | – |
| **(/Intraspecific levels)** |  |  |  |  |  |  |  |
| Within-populations (†)<br>(individuals) | 0.31 (0.25 – 0.37) | 0.30 (0.25 – 0.35) | 0.06 (0.05 – 0.06) | 0.25 (0.20 – 0.30) | 0.25 (0.20 – 0.29) | – | – |
| **Among-populations (\*)** | **0.53 (0.28 – 0.78)** | **0.49 (0.27 – 0.65)** | **0.04 (0.03 – 0.05)** | **0.49 (0.25 – 0.73)** | **0.45 (0.24 – 0.62)** | **1.80** | **.0362** |
| **(B) (/Trait types)** |  |  |  |  |  |  |  |
| Morphology (†) | 0.28 (0.22 – 0.34) | 0.27 (0.22 – 0.33) | 0.05 (0.05 – 0.06) | 0.23 (0.18 – 0.28) | 0.22 (0.18 – 0.27) | – | – |
| **Physiology (\*)** | **0.34 (0.28 – 0.41)** | **0.33 (0.27 – 0.39)** | **0.06 (0.04 – 0.07)** | **0.29 (0.24 – 0.34)** | **0.28 (0.23 – 0.33)** | **2.17** | **.0150** |
| **Behavior (\*)** | **0.33 (0.27 – 0.39)** | **0.32 (0.26 – 0.37)** | **0.06 (0.05 – 0.07)** | **0.27 (0.22 – 0.32)** | **0.26 (0.22 – 0.31)** | **1.88** | **.0301** |
| **(/Response types)** |  |  |  |  |  |  |  |
| **Foraging (\*)** | **0.38 (0.28 – 0.49)** | **0.37 (0.28 – 0.45)** | **0.07 (0.06 – 0.08)** | **0.32 (0.23 – 0.40)** | **0.31 (0.23 – 0.38)** | **2.37** | **.0178** |
| **Trophic niche (\*)** | **0.37 (0.27 – 0.48)** | **0.36 (0.26 – 0.44)** | **0.06 (0.05 – 0.07)** | **0.32 (0.22 – 0.41)** | **0.31 (0.22 – 0.39)** | **2.15** | **.0317** |
| Growth | 0.34 (0.25 – 0.43) | 0.33 (0.24 – 0.41) | 0.06 (0.05 – 0.07) | 0.28 (0.20 – 0.36) | 0.28 (0.20 – 0.35) | 1.71 | .0872 |
| Survival | 0.29 (0.20 – 0.37) | 0.28 (0.20 – 0.35) | 0.05 (0.04 – 0.06) | 0.24 (0.16 – 0.31) | 0.23 (0.16 – 0.30) | 0.94 | .3495 |
| Reproduction (†) | 0.24 (0.17 – 0.32) | 0.24 (0.16 – 0.31) | 0.05 (0.04 – 0.06) | 0.19 (0.12 – 0.26) | 0.19 (0.12 – 0.25) | – | – |
| Community | 0.39 (0.24 – 0.55) | 0.37 (0.23 – 0.50) | 0.08 (0.07 – 0.10) | 0.31 (0.17 – 0.45) | 0.30 (0.17 – 0.43) | 1.83 | .0671 |
| Ecosystem | 0.34 (0.20 – 0.48) | 0.33 (0.20 – 0.45) | 0.08 (0.07 – 0.09) | 0.26 (0.14 – 0.39) | 0.26 (0.14 – 0.37) | 0.71 | .4805 |

The intraspecific-level fixed effect (*H1*) significantly improved the fit of our model (pseudo-*R²* = 0.40; LRT= 15.13, *P* = .0005). Effect sizes estimated among populations were approximately two times larger than those at the individual level (fig. S8) and, corresponding with our main hypothesis, this difference was significant (table 2). However, *H1* was only marginally supported by a two-tailed comparison (*z* = 1.80; *P* = .0725).

The type of ecological response (*H2*) was a significant predictor of the effect sizes (pseudo-*R²* = 0.50; LRT= 20.76, *P* = .0041). Heterogeneity patterns in the individual-level dataset for the model related to *H2* were comparable to our intercept-only model fitted on the global dataset (total *I²*: 87.9%; study *I²*: 20.9%; within-study *I²*: 43.3%; phylogeny *I²*: 23.1%), but the random effect of trait type (morphology, physiology, and behavior) accounted only for a small part of the total heterogeneity (0.56%). We observed an intermediate pattern regarding our predictions for *H2*. The effect sizes for reproduction were smaller by a third than those for foraging performance and trophic niche (fig. 3, table 2). Effect sizes for community variables were 50% larger than for reproduction, but this comparison was only marginally significant (fig. 3, table 2). The same analysis carried out on the whole dataset led to qualitatively similar results, but with community responses showing significantly larger effect sizes than reproduction (table S3).

**Figure 3:**
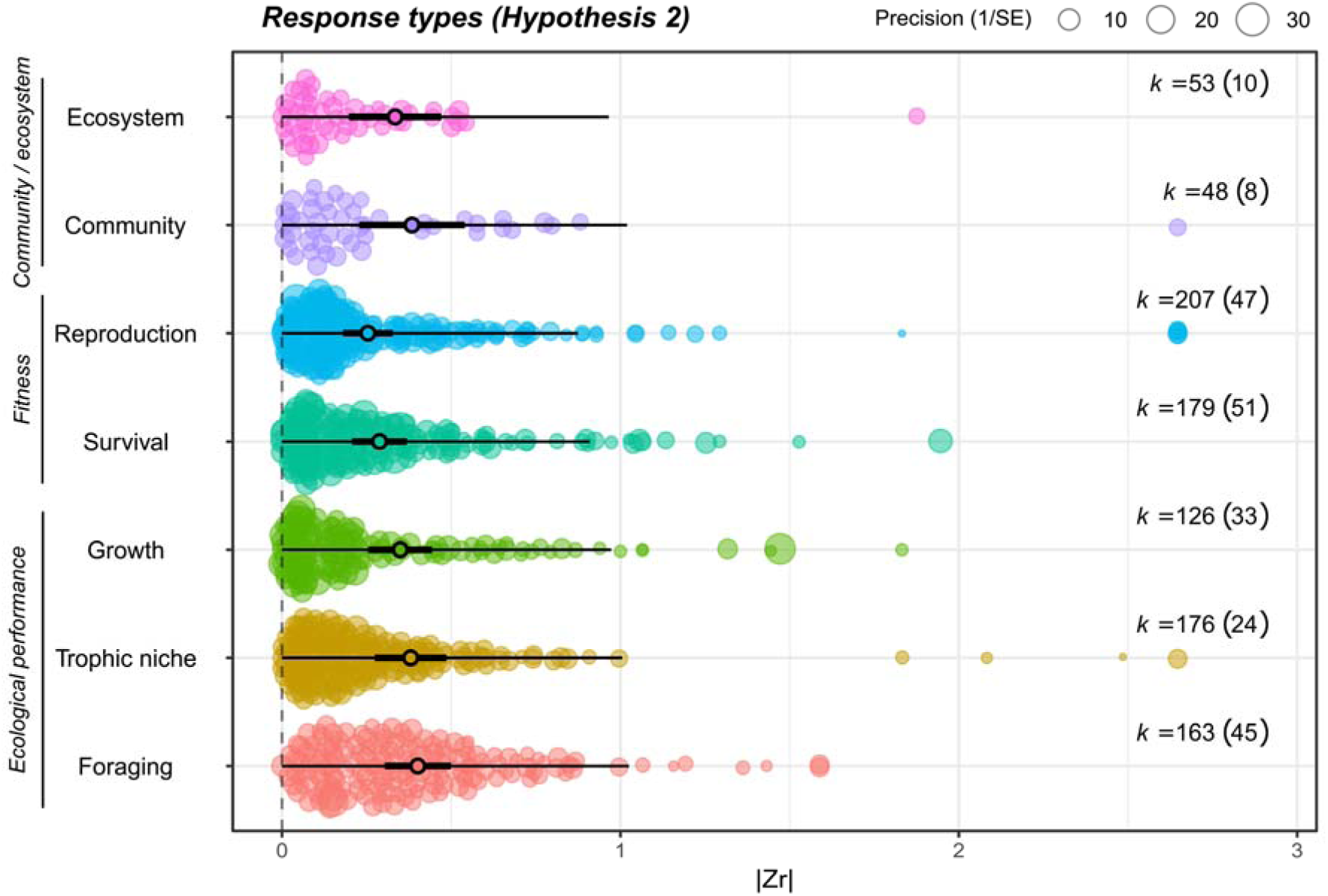
Orchard plot of model estimates of |*Zr*| effects sizes of phenotypic correlations independent from/controlling for body size, estimated for each ecological response at the individual level (n=952 effect sizes). The size of each point is proportional to the precision of the effect size (1/SE). Thick and thin error bars give 95% confidence and prediction intervals, respectively. Sample sizes (k) and number of studies (in parentheses) are given for each category of ecological responses. Model estimates are reported in table 2.

The type of phenotypic trait (*H3*) was a significant driver of the strength of relationships among the traits of individuals and ecological responses (pseudo-*R²* = 0.44; LRT= 12.05, *P* = .0072). Heterogeneity for the model for *H3* (total *I²*: 86.9%; study *I²*: 23.9%; within-study *I²*: 47.1%; phylogeny *I²*: 15.2%; response types *I²*: 0.70%) was comparable to heterogeneity in previous models (*H0-2*). In agreement with our third hypothesis (*H3)*, both physiological and behavioral traits showed effect sizes approximately 20% larger than morphological ones (fig. 4, table 2). Testing the same planned contrasts, but without any assumption of directionality showed that effect sizes for physiology were significantly larger than for morphology (two-tailed test: *z* = 2.17; *P* = .0301), whereas effect sizes for behavior were only marginally larger than for morphology (two-tailed test: *z* = 1.88; *P* = .0602). Our test of *H3* on the whole dataset (i.e., including both individual and among-population observations) showed that behavioral trait variation within species maintained stronger links to various ecological responses than morphological trait variation, but this was not the case for physiological trait variation (table S3).

**Figure 4:**
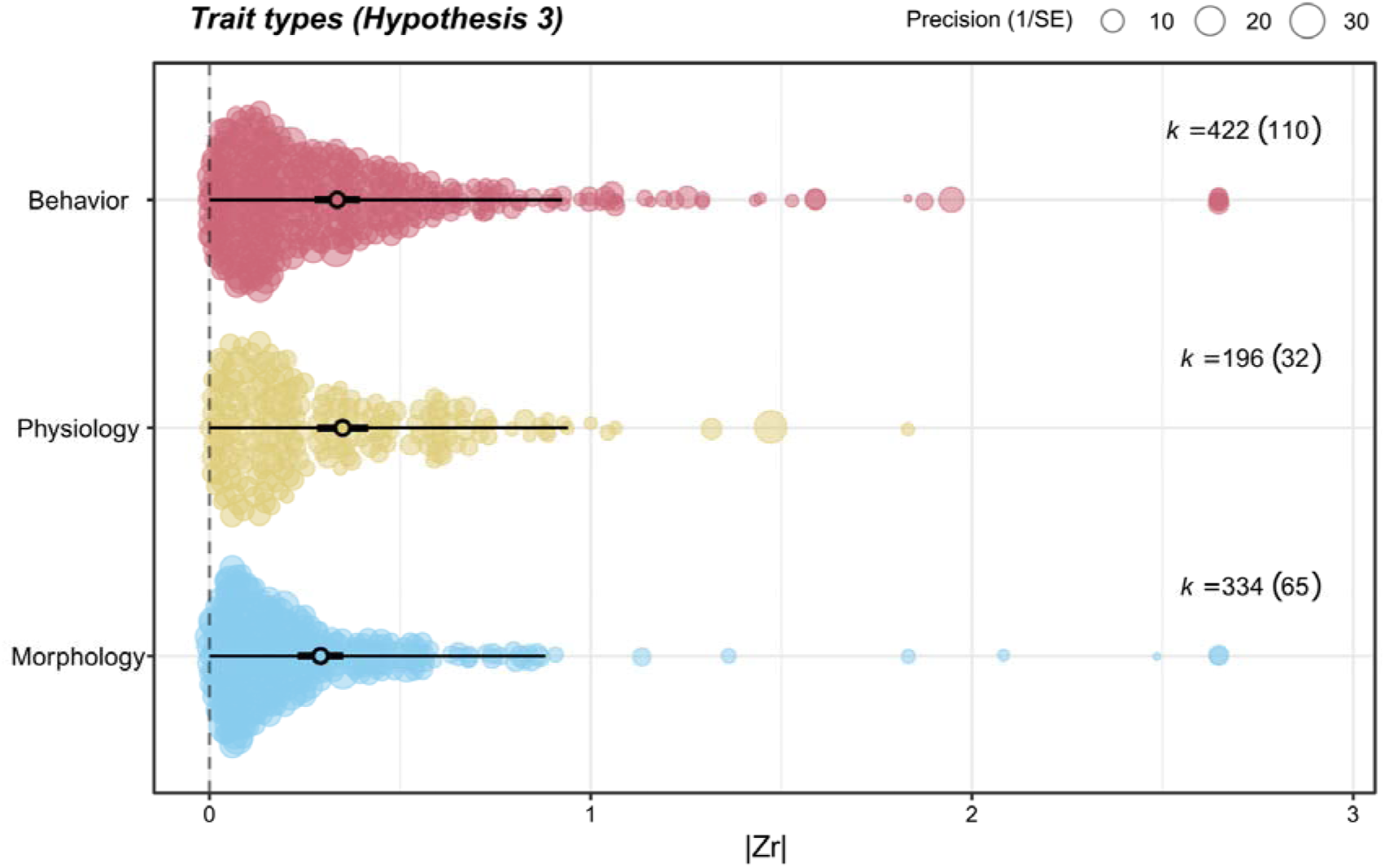
Orchard plot of model estimates of |*Zr*| effect sizes of phenotypic correlations independent from/controlling for body size, estimated for each trait type category at the individual level (n=952 effect sizes). The size of each point is proportional to the precision of the effect size (1/SE). Thick and thin error bars give 95% confidence and prediction intervals, respectively. Sample sizes (*k*) and the number of studies (in parentheses) are given for each trait type category. Model estimates are reported in table 2. Categories are shown in blue, yellow, and orange for morphological, physiological, and behavioral traits, respectively.

Immunity (n=3), hormone levels (n=11), morphotypes (n=75), and activity (n=133) were the four trait categories maintaining the strongest relationships (fig. 5). Categories with large effect sizes estimated with sufficient sample sizes were then metabolism (n=89), stoichiometry (n=52), boldness (n=91), and exploration (n=92) (fig. 5). We refrained from statistical comparisons here, as this post hoc ranking was instead aimed at a qualitative assessment of the functionality of traits in our dataset (fig. 5).

**Figure 5:**
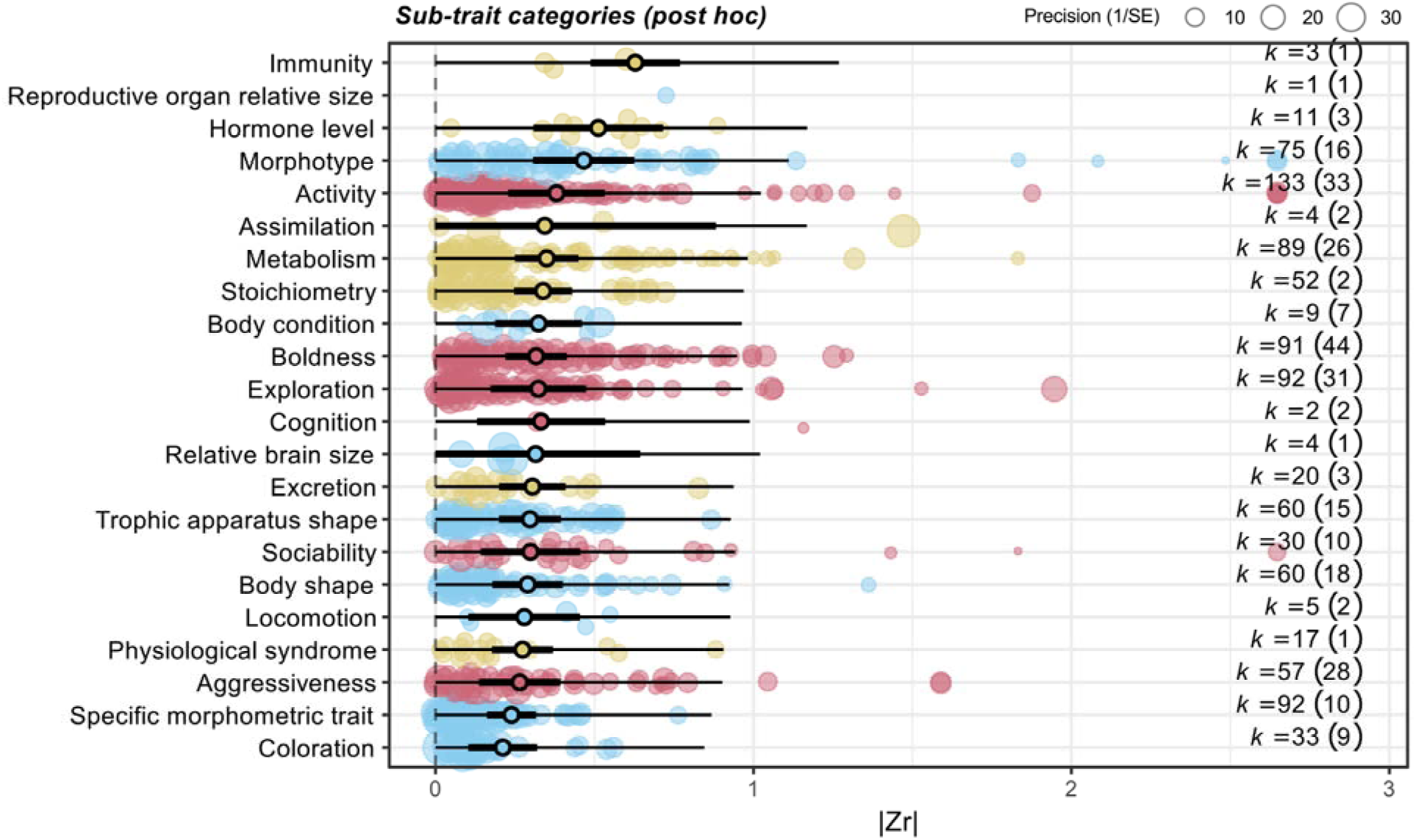
Orchard plot of model estimates of |*Zr*| effect sizes of phenotypic correlations independent from/controlling for body size, estimated for each sub-trait type category at the individual level (n=952 effect sizes), ranked from the highest (top) to the lowest estimates (bottom). The size of the bubbles is proportional to the precision of the effect sizes (1/SE). Thick and thin error bars give 95% confidence and prediction intervals, respectively. Sample sizes (*k*) and the number of studies (in parentheses) are given for each sub-trait category. Categories are shown in blue, yellow, and orange for morphological, physiological, and behavioral traits, respectively.

## Discussion

We analyzed four decades of studies in ecology and evolution to advance our understanding of the ecology of individual animals (LaBarge et al. 2024; Jeltsch et al. 2025; Szangolies et al. 2025). In providing the first broad-scale estimates of trait functionality, we emphasize that trait variability among similarly sized individuals matters to ecology, but also that the type of ecological response and the type of trait (morphology, physiology, and behavior) are significant drivers of the strength of trait-to-ecology relationships.

### Global Strength of Correlations between Intraspecific Trait Variation and Ecological Responses

Our global effect size (|*r*|_unbiased_ = 0.26 [0.21 – 0.30] 95% CI; table 2) is small to moderate in absolute terms (Nakagawa and Cuthill 2007). However, it falls in the very upper range of effect sizes reported in ecology and evolution (*ca.* 95% of |*r|* values are between 0.14 and 0.26; Møller and Jennions 2002). This remains true even when focusing solely on estimates at the individual level (|*r*|_unbiased_ = 0.25 [0.20 – 0.29] 95% CI; table 2). Surprisingly, our ad hoc analysis of the strength of correlations between individual body size and a wide range of ecological responses provided a close estimate (|*r*| = 0.28 [0.19 – 0.37] 95% CI; for a more detailed analysis, see Supplementary Materials, Section 2, fig S9 and S10). This comparison reinforces our statement that individual-level ecologies are multidimensional, and that body size alone cannot reflect all this diversity. None of the covariates we tested were significant, suggesting that our results generalize over years of publication, aquatic and terrestrial realms, consumers and predators, and methodological approaches. However, the phylogenetic signals we detected (15–23%) suggest differences across the animal kingdom in how strongly intraspecific trait variability links to ecological responses, a pattern that deserves further investigation.

Effect sizes at the level of populations and colonies of eusocial animals were large (|*r*|_unbiased_ = 0.45 [0.24 – 0.62] 95% CI; table 2), nearly two times larger than at the individual level (fig. S8). This contrast supports our hypothesis *H1* that populations that diverged phenotypically would show strong links between traits and ecological responses. In a world where genetic and phenotypic diversity is changing rapidly (Exposito-Alonso et al. 2022; Sanderson et al. 2023), maintaining high population-level phenotypic diversity matters for the conservation and management of ecosystems (Blanchet et al. 2020; Des Roches et al. 2021).

### Strength of Correlations between Individual Trait Variation and Ecological Responses Differs Across Biological Levels

Due to the potential accumulation of intermediate relationships between the traits of an individual and distal responses (see Bailey et al. 2009), we expected (*H2*) that the overall strength of relationships would decrease from proximal responses (low biological level), such as food intake or growth rate, to distal responses at higher levels of biological organization (e.g., communities and ecosystems). We instead observed a mixed pattern, with moderately high correlations for both ecological performance and community responses, whereas fitness (survival and reproduction) displayed the smallest effect sizes. Reproductive outcomes are complex, as they are influenced by intrinsic (e.g., genetic) and extrinsic processes such as changing environments and random events occurring over the reproductive life of an animal (Merilä and Sheldon 2000), which can reduce the effect sizes we reported for reproduction. Large effect sizes for community responses could arise because of greater complexity in nature than under controlled conditions (Hendry 2019). Most of the studies investigating fitness took place in nature (*ca.* 62%), whereas most studies investigating community and ecosystem responses were conducted in mesocosms (*ca.* 87%). However, we did not find a difference among field, mesocosm and microcosm effect sizes. Although further investigation is needed in natural settings, we are left with the result that correlations between individual trait variability and community dynamics can be strong (Des Roches et al. 2018; Raffard et al. 2019), even when they are maintained by phenotypic differences among similarly sized individuals. Interestingly, our analysis of an *ad hoc* dataset of selected meta-analysis and empirical studies reporting relationships between body size variation and various ecological responses showed a similar pattern, namely that the smallest effect sizes were for fitness, whereas foraging and community/ecosystem responses exhibited larger effect sizes (Supplementary Materials, Section 2, fig. S9 and S10).

The estimates for fitness proxies reported here (table 2) are of similar nature to linear estimates of directional selection (Hendry 2017). Indeed, the strength of correlations we estimated between traits and reproduction (|*r*|_unbiased_ = 0.19) matches previous estimates of directional selection (|β| = 0.16; Kingsolver et al. 2001; Kingsolver et al. 2012). One reason for the relatively small effect sizes observed for fitness could be that adaptation already selected traits near their optimum, reducing the number of phenotypes that deviate far from optimum. As such, estimated selection is no longer strong because “selection erases its traces” (Haller and Hendry 2014). Another mechanism can be fluctuating selection, which averages out selection signals over time (Wright et al. 2019). The links between trait variability and community assembly, however, may not reflect adaptations *per se*, and are probably not under selection to reduce such effects, which may explain why effect sizes for community responses tended to be larger than those for reproduction.

A third argument comes from behavioral ecology. Labile traits such as behavior and physiology harbor a reversible or contextual trait variance component (which is less the case for structural, morphological traits), reflecting variation in the expression of a trait for a given individual across situations and time (i.e., intra-individual trait variability); along with an among-individual trait variation component, which corresponds to the consistent component of trait expression for a given individual across time or contexts (Sih et al. 2004; Réale et al. 2007). A poor partitioning of trait variance among these two components can dampen relationships (Niemelä and Dingemanse 2018*b*). In re-examining our estimates of fitness correlates (386 effect sizes from 90 studies assessing survival and reproduction), we find that physiology, and, in a lesser extent, behavior, exceed morphological traits in terms of correlation strengths with fitness (one-tailed planned contrasts: *z* = 2.12; *P* = .0171 and *z* = 1.39; *P* = .0829, for physiology and behavior, respectively), thereby providing only limited support for this rationale.

Many relationships in ecology are non-linear (Bolnick et al. 2011). For instance, due to Jensen’s inequality, the accumulation of concave-up intermediate relationships can magnify the effects of trait variability at high levels of biological organization (see Bailey et al. 2009). In addition, large effects can occur through direct effects on community and ecosystem responses (i.e., ‘trait-mediated effects’; Schmitz et al. 2004; see fig. 1a). This mechanism may be particularly relevant when focal animals are keystone (Paine 1980), engineers (Romero et al. 2015), or prone to niche construction (Laland et al. 1999). A great example comes from the Grand Voyageur National Park. A small subset of wolves within a pack specialized in hunting twice as many beavers as others (Bump et al. 2022), raising the question of whether behavioral differences underlie this hunting specialization by keystone individuals that disproportionately reshapes wetlands (Gable et al. 2020).

### Do Trait Types Show Different Strengths of Relationships with Ecological Responses?

We hypothesized that differences in physiology and behavior among individuals would generate stronger ecological relationships than would differences in morphology (*H3*). Consequently, physiology and behavior showed effects sizes 20% larger than those for morphology. These differences, albeit modest on average, as marked by highly variable effect sizes within each trait class, still support the idea that traits reflecting energetic currencies and “what individuals do” underlie ecological performance, fitness, interactions among individuals, and therefore drive community and ecosystem processes (Szangolies et al. 2025) – even as much as does body size variation among individuals (Supplementary Materials, Section 2). For example, functional responses of predator crabs (*Panopeus herbstii*) on mussel prey (*Brachidontes exustus*) are mediated by consistent variation in activity levels among individuals (Toscano and Griffen 2014), whose effects may cascade at the ecosystem level since filter-feeding mussels are ecosystem engineers. Similarly, physiological traits showed large effect sizes across ecological responses (fig. 4 and 5) and, as our post hoc analysis in support of the discussion showed above, this difference seems particularly true in the case of fitness. A good example is that variation in corticosterone levels predicts long-term reproductive performance among wandering albatross individuals (*Diomedea exulans*; Angelier et al. 2010). Metabolic, immune, hormonal, and stoichiometric traits have all mechanistic links with fitness, community, and ecosystem responses (Elser et al. 2000; Ricklefs and Wikelski 2002; Brandl et al. 2022), and should be an increasing focus in the trait-based ecology of animals.

Understanding why, fundamentally, different phenotypic dimensions of an individual may vary in their correlation strengths with ecology is challenging. One may see morphology as a *potential* for the ecological effects of a phenotype, whereas physiology and behavior might be closer to *realized* effects (Wainwright and Reilly 1994). Our results agree with this view, although the average differences of effect sizes among those trait types were rather small and variation was extremely high within trait classes (figures 4 and 5; table 2). However, our detailed analysis of trait categories showed perhaps not surprisingly that morphological traits with the largest effect sizes had clear functional interpretation. ‘*Morphotypes*’ are dimorphic ecotypes which may have diverged hundreds to thousands of generations ago (Harmon et al. 2009). ‘*Body condition*’ is a proxy for physiological status (energy reserve). However, other traits with specific functional relevance, such as specific linear morphometric traits (e.g., relative tarsus length), exhibited lower effect sizes. Perhaps the safest interpretation here is that different types of morphological traits could show very different strengths of effects on ecology, with some being very large, and others very small, yet morphology as a trait class showed low effect sizes on average.

### Concluding Remarks

Our results at the individual level emphasizing physiology and behavior are somewhat contrasting with the strong reliance on morphology (and body size) in trait-based ecology (Dawson et al. 2021). The popularity of morphology may lie in its prominent role in the early development of the niche variation hypothesis (Van Valen 1965) – or simply because morphological traits are easy to measure. Another reason for its popularity could be its low intra-individual variation and high heritability compared to physiology and behavior (Mousseau and Roff 1987). Although morphology shows weaker ecological associations in our analyses, one could argue that its greater repeatability and heritability relative to physiology and behavior may promote stronger eco-evolutionary dynamics. However, the lability of physiology and behavior does not imply that this variation is random, and the perceived low heritability of behavior does not necessarily indicate low additive genetic variance or evolvability (for detailed explanations, see Houle 1992; Hansen et al. 2011). In fact, physiology and behavior often show moderate to high repeatability among individuals (Bell et al. 2009) and, in some cases, large additive genetic variance and heritabilities (Dochtermann et al. 2014). Moreover, intra-individual variation in behavior can be structured along a gradient from predictable to unpredictable individuals (Stamps et al. 2012), yet the ecological implications of this are largely unknown. To assess the propensity of a broad range of traits to be involved in eco-evolutionary dynamics, it would be promising to compare their ecological functionality (as we report here) to evolutionary parameters such as evolvability (Hansen et al. 2011). Another avenue of interest would be to better understand the phenotypic determinants of niche construction and nongenetic (including ecological) inheritance (Laland et al. 1999; Danchin et al. 2011). These joint estimates could then be used to parameterize eco-evolutionary models that assess how shifts in trait distributions, or the emergence of behavioral innovations, affect the evolution of characters, ecological dynamics, and their feedbacks (Govaert et al. 2019). Considering future eco-evolutionary syntheses compiling information on a wide array of trait types, we could imagine testing the prevalence of trade-offs across traits – for instance, traits showing large evolvability but weak ecological effects, or, conversely, traits with both high evolvability and strong ecological effects. Such future syntheses can advance us toward a unified understanding of the ecological and evolutionary processes associated with different types of biological attributes.

To unravel the consequences of individual trait variation in animals, we encourage ecologists to pay greater attention to physiology and behavior, in nature, especially when investigating their effects on community and ecosystem levels. A step forward would be to better acknowledge the implications of trait types and syndromes, their ecological tradeoffs, and their consequences on ecosystem processes. Future studies will gain improved predictive abilities by simultaneously addressing all the entangled facets of a phenotype (Arnold 1983; Violle et al. 2007). Such an approach to the ecological consequences of individual trait (co)variations will complement the synthesis we provide here, by improving the integration of biological scales with evolutionary and ecological processes, as well as enhancing our ability to predict the effects of human-driven changes on rapidly evolving ecosystems (Blanchet et al. 2020; Des Roches et al. 2021).

## Acknowledgments

We are grateful to the researchers who contributed, from early discussions, to our thinking of the issue, especially J. Cucherousset and L. Závorka. We thank B. Corbara for the discussion about our results. We also thank A. Bruder, R. Oester, A. M. Roth, L. Govaert, T. Norin, C. Crafter, and J. O’Brien for their suggestions on the manuscript before its submission to the journal. We are thankful to the editors and anonymous reviewers, whose comments greatly improved the final work. T. Rota thanks the French Agence Nationale de la Recherche (ANR) for its support through a grant called ‘RESILIENCE’ [ANR-18-CE02-0015], the Swiss National Foundation for Science (SNFS) for a grant called ‘MULTIBEF’ [315230_204998], and the Czech Science Foundation (GAČR), for a grant called ‘POLLCHAIN’ [project No. 25-17362S]. Financial support was provided to R. Céréghino by the French Agence Nationale de la Recherche (ANR) through an Investissement d’Avenir grant [LabEx CEBA, ref. ANR-10-LABX-25-01]. The authors have no conflict of interest to declare.

## Statement of Authorship

T.R. assembled the first ideas. T.R. designed the study with the guidance of A.R., S.B., A.P.H., and R.C.. T.R. and A.R. performed the systematic review of the literature in parallel, deleted duplicates and validated selected studies. T.R. extracted the data from the selected papers, designed and performed the statistical analyses, wrote a first draft of the manuscript, and edited it until its final form. All co-authors contributed to the editing of the manuscript until its final form.

## Data and Code Availability

The data and R code reproducing the results can be accessed through a public and permanent repository (Figshare), at https://doi.org/10.6084/m9.figshare.27377529.

## Literature Cited

Albert, C. H. 2015. Intraspecific trait variability matters. Journal of Vegetation Science 26:7–8.

Alberti, M., C. Correa, J. M. Marzluff, A. P. Hendry, E. P. Palkovacs, K. M. Gotanda, V. M. Hunt, et al. 2017. Global urban signatures of phenotypic change in animal and plant populations. Proceedings of the National Academy of Sciences 114:8951–8956.

Angelier, F., J. C. Wingfield, H. Weimerskirch, and O. Chastel. 2010. Hormonal correlates of individual quality in a long-lived bird: a test of the ‘corticosterone–fitness hypothesis.’ Biology Letters 6:846–849.

Arnold, S. J. 1983. Morphology, Performance and Fitness. American Zoologist 23:347–361.

Bailey, J. K., J. A. Schweitzer, F. Ubeda, J. Koricheva, C. J. LeRoy, M. D. Madritch, B. J. Rehill, et al. 2009. From genes to ecosystems: a synthesis of the effects of plant genetic factors across levels of organization. Philosophical Transactions of the Royal Society B: Biological Sciences 364:1607–1616.

Baldwin, J. M. 1897. Organic Selection. Nature 55:558–558.

Barabás, G., C. Parent, A. Kraemer, F. Van de Perre, and F. De Laender. 2022. The evolution of trait variance creates a tension between species diversity and functional diversity. Nature Communications 13:2521.

Bell, A. M., S. J. Hankison, and K. L. Laskowski. 2009. The repeatability of behaviour: a meta-analysis. Animal Behaviour 77:771–783.

Biro, P. A., and J. A. Stamps. 2008. Are animal personality traits linked to life-history productivity? Trends in Ecology & Evolution 23:361–368.

Blanchet, S., J. G. Prunier, I. Paz-Vinas, K. Saint-Pé, O. Rey, A. Raffard, E. Mathieu-Bégné, et al. 2020. A river runs through it: The causes, consequences, and management of intraspecific diversity in river networks. Evolutionary Applications 13:1195–1213.

Boag, P. T., and P. R. Grant. 1981. Intense Natural Selection in a Population of Darwin’s Finches (Geospizinae) in the Galapagos. Science 214:82–85.

Bolnick, D. I., P. Amarasekare, M. S. Araújo, R. Bürger, J. M. Levine, M. Novak, V. H. W. Rudolf, et al. 2011. Why intraspecific trait variation matters in community ecology. Trends in Ecology & Evolution 26:183–192.

Bolnick, D. I., R. Svanbäck, M. S. Araújo, and L. Persson. 2007. Comparative support for the niche variation hypothesis that more generalized populations also are more heterogeneous. Proceedings of the National Academy of Sciences 104:10075–10079.

Brandl, S. J., J. S. Lefcheck, A. E. Bates, D. B. Rasher, and T. Norin. 2022. Can metabolic traits explain animal community assembly and functioning? Biological Reviews brv.12892.

Brose, U., T. Jonsson, E. L. Berlow, P. Warren, C. Banasek-Richter, L.-F. Bersier, J. L. Blanchard, et al. 2006. Consumer–resource body-size relationships in natural food webs. Ecology 87:2411–2417.

Brown, J. H., J. F. Gillooly, A. P. Allen, V. M. Savage, and G. B. West. 2004. Toward a metabolic theory of ecology. Ecology 85:1771–1789.

Bump, J., T. Gable, S. Johnson-Bice, A. Homkes, D. Freund, S. Windels, and S. Chakrabarti. 2022. Predator personalities alter ecosystem services. Frontiers in Ecology and the Environment 20:275–277.

Careau, V., and T. Garland. 2012. Performance, Personality, and Energetics: Correlation, Causation, and Mechanism. Physiological and Biochemical Zoology 85:543–571.

Careau, V., S. S. Killen, and N. B. Metcalfe. 2014. Adding Fuel to the “Fire of Life”: Energy Budgets across Levels of Variation in Ectotherms and Endotherms, Integrative Oganismal Biology (eds), John Wiley & Sons, Inc, Hoboken, NJ, USA. doi: 10.1002/9781118398814.ch14. Pages 219–234 in Integrative Oganismal Biology (John Wiley&Sons.). L. B. Martin, C. K. Ghalambor and H. A. Woods, Hoboken, NJ, USA.

Careau, V., D. Thomas, M. M. Humphries, and D. Réale. 2008. Energy metabolism and animal personality. Oikos 117:641–653.

Carmona, C. P., F. de Bello, N. W. H. Mason, and J. Lepš. 2016. Traits Without Borders: Integrating Functional Diversity Across Scales. Trends in Ecology & Evolution 31:382–394.

Ceballos, G., P. R. Ehrlich, and R. Dirzo. 2017. Biological annihilation via the ongoing sixth mass extinction signaled by vertebrate population losses and declines. Proceedings of the National Academy of Sciences 114.

Chacón-Labella, J., C. Hinojo-Hinojo, T. Bohner, M. Castorena, C. Violle, V. Vandvik, and B. J. Enquist. 2022. How to improve scaling from traits to ecosystem processes. Trends in Ecology & Evolution S0169534722002749.

Cianciaruso, M. V., M. A. Batalha, K. J. Gaston, and O. L. Petchey. 2009. Including intraspecific variability in functional diversity. Ecology 90:81–89.

Danchin, É., A. Charmantier, F. A. Champagne, A. Mesoudi, B. Pujol, and S. Blanchet. 2011. Beyond DNA: integrating inclusive inheritance into an extended theory of evolution. Nature Reviews Genetics 12:475–486.

Darwin, C., and A. Wallace. 1858. On the Tendency of Species to form Varieties; and on the Perpetuation of Varieties and Species by Natural Means of Selection. Journal of the Proceedings of the Linnean Society of London. Zoology 3:45–62.

Dawson, S. K., C. P. Carmona, M. González-Suárez, M. Jönsson, F. Chichorro, M. Mallen-Cooper, Y. Melero, et al. 2021. The traits of “trait ecologists”: An analysis of the use of trait and functional trait terminology. Ecology and Evolution 11:16434–16445.

de Bello, F., Z. Botta-Dukát, J. Lepš, and P. Fibich. 2021. Towards a more balanced combination of multiple traits when computing functional differences between species. Methods in Ecology and Evolution 12:443–448.

Des Roches, S., L. H. Pendleton, B. Shapiro, and E. P. Palkovacs. 2021. Conserving intraspecific variation for nature’s contributions to people. Nature Ecology & Evolution 5:574–582.

Des Roches, S., D. M. Post, N. E. Turley, J. K. Bailey, A. P. Hendry, M. T. Kinnison, J. A. Schweitzer, et al. 2018. The ecological importance of intraspecific variation. Nature Ecology & Evolution 2:57–64.

Des Roches, S., J. B. Shurin, D. Schluter, and L. J. Harmon. 2013. Ecological and Evolutionary Effects of Stickleback on Community Structure. PLoS ONE 8:e59644.

Dochtermann, N. A., T. Schwab, and A. Sih. 2014. The contribution of additive genetic variation to personality variation: heritability of personality. Proceedings of the Royal Society B: Biological Sciences 282:20142201–20142201.

Elser, J. j., R. w. Sterner, E. Gorokhova, W. f. Fagan, T. a. Markow, J. b. Cotner, J. f. Harrison, et al. 2000. Biological stoichiometry from genes to ecosystems. Ecology Letters 3:540–550.

Elton, C. S. 1927. Animal ecology, by Charles Elton; with an introduction by Julian S. Huxley. Macmillan Co., New York.

Enquist, B. J., J. Norberg, S. P. Bonser, C. Violle, C. T. Webb, A. Henderson, L. L. Sloat, et al. 2015. Scaling from Traits to Ecosystems. Pages 249–318 in Advances in Ecological Research (Vol. 52). Elsevier.

Exposito-Alonso, M., T. R. Booker, L. Czech, L. Gillespie, S. Hateley, C. C. Kyriazis, P. L. M. Lang, et al. 2022. Genetic diversity loss in the Anthropocene. Science 377:1431–1435.

Funk, J. L., J. E. Larson, G. M. Ames, B. J. Butterfield, J. Cavender-Bares, J. Firn, D. C. Laughlin, et al. 2017. Revisiting the H oly G rail: using plant functional traits to understand ecological processes. Biological Reviews 92:1156–1173.

Gable, T. D., S. M. Johnson-Bice, A. T. Homkes, S. K. Windels, and J. K. Bump. 2020. Outsized effect of predation: Wolves alter wetland creation and recolonization by killing ecosystem engineers. Science Advances 6:eabc5439.

Gibert, J. P., A. I. Dell, J. P. DeLong, and S. Pawar. 2015. Scaling-up Trait Variation from Individuals to Ecosystems. Pages 1–17 in Advances in Ecological Research (Vol. 52). Elsevier.

Gifford, M. E., T. A. Clay, and V. Careau. 2014. Individual (Co)variation in Standard Metabolic Rate, Feeding Rate, and Exploratory Behavior in Wild-Caught Semiaquatic Salamanders. Physiological and Biochemical Zoology 87:384–396.

Gordon, D. M. 2011. The fusion of behavioral ecology and ecology. Behavioral Ecology 22:225–230.

Govaert, L., E. A. Fronhofer, S. Lion, C. Eizaguirre, D. Bonte, M. Egas, A. P. Hendry, et al. 2019. Eco-evolutionary feedbacks—Theoretical models and perspectives. Functional Ecology 33:13–30.

Grafen, A. 1989. The phylogenetic regression. Philosophical Transactions of the Royal Society of London. B, Biological Sciences 326:119–157.

Green, S. J., C. B. Brookson, N. A. Hardy, and L. B. Crowder. 2022. Trait-based approaches to global change ecology: moving from description to prediction. Proceedings of the Royal Society B: Biological Sciences 289:20220071.

Hall, B. K. 2001. Organic Selection: Proximate Environmental Effects on the Evolution of Morphology and Behaviour. Biology and Philosophy 16:215–237.

Haller, B. C., and A. P. Hendry. 2014. Solving the paradox of stasis: squashed stabilizing selection and the limits of detection: selection and the limits of detection. Evolution 68:483–500.

Hansen, T. F., C. Pélabon, and D. Houle. 2011. Heritability is not Evolvability. Evolutionary Biology 38:258–277.

Harmon, L. J., B. Matthews, S. Des Roches, J. M. Chase, J. B. Shurin, and D. Schluter. 2009. Evolutionary diversification in stickleback affects ecosystem functioning. Nature 458:1167–1170.

Hart, S. P., S. J. Schreiber, and J. M. Levine. 2016. How variation between individuals affects species coexistence. Ecology Letters 19:825–838.

Hendry, A. P. 2017. Eco-evolutionary dynamics. Princeton University Press, Princeton.

Hendry, A. P. 2019. A critique for eco-evolutionary dynamics. Functional Ecology 33:84–94.

Hinchliff, C. E., S. A. Smith, J. F. Allman, J. G. Burleigh, R. Chaudhary, L. M. Coghill, K. A. Crandall, et al. 2015. Synthesis of phylogeny and taxonomy into a comprehensive tree of life. Proceedings of the National Academy of Sciences 112:12764–12769.

Hothorn, T., F. Bretz, and P. Westfall. 2008. Simultaneous Inference in General Parametric Models. Biometrical Journal 50:346–363.

Houle, D. 1992. Comparing evolvability and variability of quantitative traits. Genetics 130:195–204.

Hutchinson, G. E. 1957. Concluding Remarks. Cold Spring Harbor Symposia on Quantitative Biology 22:415–427.

Jeltsch, F., M. Roeleke, A. Abdelfattah, R. Arlinghaus, G. Berg, N. Blaum, L. De Meester, et al. 2025. The need for an individual-based global change ecology. Individual-based Ecology 1:1–18.

Kingsolver, J. G., S. E. Diamond, A. M. Siepielski, and S. M. Carlson. 2012. Synthetic analyses of phenotypic selection in natural populations: lessons, limitations and future directions. Evolutionary Ecology 26:1101–1118.

Kingsolver, J. G., and S. E. Diamond. 2011. Phenotypic Selection in Natural Populations: What Limits Directional Selection? The American Naturalist 177:346–357.

Kingsolver, J. G., H. E. Hoekstra, J. M. Hoekstra, D. Berrigan, S. N. Vignieri, C. E. Hill, A. Hoang, et al. 2001. The Strength of Phenotypic Selection in Natural Populations. The American Naturalist 157:245–261.

Kinnison, M. T., and A. P. Hendry. 2001. The pace of modern life II: from rates of contemporary microevolution to pattern and process. Genetica 112–113:145–164.

Kleiber, M. 1932. Body size and metabolism. Hilgardia 6:315–353.

LaBarge, L. R., M. Krofel, M. L. Allen, R. A. Hill, A. J. Welch, and A. T. L. Allan. 2024. Keystone individuals – linking predator traits to community ecology. Trends in Ecology & Evolution 39:983–994.

Laland, K. N. 2004. Extending the Extended Phenotype. Biology & Philosophy 19:313–325.

Laland, K. N., F. J. Odling-Smee, and M. W. Feldman. 1999. Evolutionary consequences of niche construction and their implications for ecology. Proceedings of the National Academy of Sciences 96:10242–10247.

Lang, I., I. Paz-Vinas, J. Cucherousset, and G. Loot. 2021. Patterns and determinants of phenotypic variability within two invasive crayfish species. Freshwater Biology 66:1782–1798.

Lawton, J. H. 1994. What Do Species Do in Ecosystems? Oikos 71:367.

Lecerf, A., and E. Chauvet. 2008. Intraspecific variability in leaf traits strongly affects alder leaf decomposition in a stream. Basic and Applied Ecology 9:598–605.

Lefcheck, J. S., V. A. G. Bastazini, and J. N. Griffin. 2015. Choosing and using multiple traits in functional diversity research. Environmental Conservation 42:104–107.

Macarthur, R., and R. Levins. 1967. The Limiting Similarity, Convergence, and Divergence of Coexisting Species. The American Naturalist 101:377–385.

McCollum, S. A., and J. Van Buskirk. 1996. Costs and benefits of a predator-induced polyphenism in the gray treefrog *Hyla chrysoscelis*. Evolution 50:583–593.

McGhee, K. E., L. M. Pintor, and A. M. Bell. 2013. Reciprocal Behavioral Plasticity and Behavioral Types during Predator-Prey Interactions. The American Naturalist 182:704–717.

McGill, B., B. Enquist, E. Weiher, and M. Westoby. 2006. Rebuilding community ecology from functional traits. Trends in Ecology & Evolution 21:178–185.

Merilä, J., and B. C. Sheldon. 2000. Lifetime Reproductive Success and Heritability in Nature. The American Naturalist 155:301–310.

Messier, J., B. J. McGill, and M. J. Lechowicz. 2010. How do traits vary across ecological scales? A case for trait-based ecology: How do traits vary across ecological scales? Ecology Letters 13:838–848.

Michonneau, F., J. W. Brown, and D. J. Winter. 2016. rotl: an R package to interact with the Open Tree of Life data. Methods in Ecology and Evolution 7:1476–1481.

Mlambo, M. C. 2014. Not all traits are ‘functional’: insights from taxonomy and biodiversity-ecosystem functioning research. Biodiversity and Conservation 23:781–790.

Moiron, M., K. L. Laskowski, and P. T. Niemelä. 2020. Individual differences in behaviour explain variation in survival: a meta-analysis. Ecology Letters 23:399–408.

Møller, A., and M. D. Jennions. 2002. How much variance can be explained by ecologists and evolutionary biologists? Oecologia 132:492–500.

Moran, N. P., A. Sánchez-Tójar, H. Schielzeth, and K. Reinhold. 2021. Poor nutritional condition promotes high-risk behaviours: a systematic review and meta-analysis. Biological Reviews 96:269–288.

Moran, N. P., B. B. M. Wong, and R. M. Thompson. 2017. Weaving animal temperament into food webs: implications for biodiversity. Oikos 126:917–930.

Morrissey, M. B. 2016. Meta-analysis of magnitudes, differences and variation in evolutionary parameters. Journal of Evolutionary Biology 29:1882–1904.

Mousseau, T. A., and D. A. Roff. 1987. Natural selection and the heritability of fitness components. Heredity 59:181–197.

Nakagawa, S., and I. C. Cuthill. 2007. Effect size, confidence interval and statistical significance: a practical guide for biologists. Biological Reviews 82:591–605.

Nakagawa, S., M. Lagisz, M. D. Jennions, J. Koricheva, D. W. A. Noble, T. H. Parker, A. Sánchez-Tójar, et al. 2022. Methods for testing publication bias in ecological and evolutionary meta-analyses. Methods in Ecology and Evolution 13:4–21.

Nakagawa, S., M. Lagisz, R. E. O’Dea, P. Pottier, J. Rutkowska, A. M. Senior, Y. Yang, et al. 2023. orchaRd 2.0: An R package for visualising meta-analyses with orchard plots. Methods in Ecology and Evolution 14:2003–2010.

Niemelä, P. T., and N. J. Dingemanse. 2018a. Meta-analysis reveals weak associations between intrinsic state and personality. Proceedings of the Royal Society B: Biological Sciences 285:20172823.

Niemelä, P. T., and N. J. Dingemanse. 2018b. On the usage of single measurements in behavioural ecology research on individual differences. Animal Behaviour 145:99–105.

Noble, D. W. A., M. Lagisz, R. E. O’dea, and S. Nakagawa. 2017. Nonindependence and sensitivity analyses in ecological and evolutionary meta-analyses. Molecular Ecology 26:2410–2425.

Nyqvist, M. J., J. Cucherousset, R. E. Gozlan, and J. R. Britton. 2018. Relationships between individual movement, trophic position and growth of juvenile pike (*Esox lucius)*. Ecology of Freshwater Fish 27:398–407.

O’Dea, R. E., M. Lagisz, M. D. Jennions, J. Koricheva, D. W. A. Noble, T. H. Parker, J. Gurevitch, et al. 2021. Preferred reporting items for systematic reviews and meta-analyses in ecology and evolutionary biology: a PRISMA extension. Biological Reviews 96:1695–1722.

Paine, R. T. 1980. Food Webs: Linkage, Interaction Strength and Community Infrastructure. The Journal of Animal Ecology 49:666.

Paradis, E., and K. Schliep. 2019. ape 5.0: an environment for modern phylogenetics and evolutionary analyses in R. Bioinformatics 35:526–528.

Pelletier, F., D. Garant, and A. P. Hendry. 2009. Eco-evolutionary dynamics. Philosophical Transactions of the Royal Society B: Biological Sciences 364:1483–1489.

Petchey, O. L., and K. J. Gaston. 2006. Functional diversity: back to basics and looking forward. Ecology Letters 9:741–758.

Peters, R. H. 1993. The ecological implications of body size. Univ. Press, Cambridge.

Pianka, E. R. 1970. On r- and K-Selection. The American Naturalist 104:592–597.

Post, D. M., E. P. Palkovacs, E. G. Schielke, and S. I. Dodson. 2008. Intraspecific variation in a predator affects community structure and cascading trophic interactions. Ecology 89:2019–2032.

Raffard, A., J. Cucherousset, J. M. Montoya, M. Richard, S. Acoca-Pidolle, C. Poésy, A. Garreau, et al. 2021. Intraspecific diversity loss in a predator species alters prey community structure and ecosystem functions. PLOS Biology 19:e3001145.

Raffard, A., J. Cucherousset, F. Santoul, L. Di Gesu, and S. Blanchet. 2023. Climate and intraspecific variation in a consumer species drive ecosystem multifunctionality. Oikos 2023:e09286.

Raffard, A., A. Lecerf, J. Cote, M. Buoro, R. Lassus, and J. Cucherousset. 2017. The functional syndrome: linking individual trait variability to ecosystem functioning. Proceedings of the Royal Society B: Biological Sciences 284:20171893.

Raffard, A., F. Santoul, J. Cucherousset, and S. Blanchet. 2019. The community and ecosystem consequences of intraspecific diversity: a meta-analysis: The ecological effects of intraspecific diversity. Biological Reviews.

Réale, D., D. Garant, M. M. Humphries, P. Bergeron, V. Careau, and P.-O. Montiglio. 2010. Personality and the emergence of the pace-of-life syndrome concept at the population level. Philosophical Transactions of the Royal Society B: Biological Sciences 365:4051–4063.

Réale, D., S. M. Reader, D. Sol, P. T. McDougall, and N. J. Dingemanse. 2007. Integrating animal temperament within ecology and evolution. Biological Reviews 82:291–318.

Ricklefs, R. E., and M. Wikelski. 2002. The physiology/life-history nexus. Trends in Ecology & Evolution 17:462–468.

Romero, G. Q., T. Gonçalves-Souza, C. Vieira, and J. Koricheva. 2015. Ecosystem engineering effects on species diversity across ecosystems: a meta-analysis: Ecosystem engineering effects across ecosystems. Biological Reviews 90:877–890.

Rota, T., J. Jabiol, E. Chauvet, and A. Lecerf. 2018. Phenotypic determinants of inter-individual variability of litter consumption rate in a detritivore population. Oikos 127:1670–1678.

Rota, T., A. Lecerf, É. Chauvet, and B. Pey. 2022. The importance of intraspecific variation in litter consumption rate of aquatic and terrestrial macro-detritivores. Basic and Applied Ecology 63:175–185.

Rota, T., J. Jabiol, S. Lamothe, D. Lambrigot, É. Chauvet, and A. Lecerf. 2024. Phenotypes of predator individuals underpin contrasting ecosystem effects. Freshwater Biology 69: 739–753.

Rota, T. 2026. Data and code from: [Behavior and Physiology Outpace Form When Linking Traits to Ecological Responses Within Populations: A Meta-Analysis]. American Naturalist, Figshare, Version 5, 10.6084/m9.figshare.27377529.

Sanderson, S., D. I. Bolnick, M. T. Kinnison, R. E. O’Dea, L. D. Gorné, A. P. Hendry, and K. M. Gotanda. 2023. Contemporary changes in phenotypic variation, and the potential consequences for eco-evolutionary dynamics. Ecology Letters 26.

Schleuning, M., D. García, and J. A. Tobias. 2023. Animal functional traits: Towards a trait-based ecology for whole ecosystems. Functional Ecology 37:4–12.

Schmitz, O. J., V. Krivan, and O. Ovadia. 2004. Trophic cascades: the primacy of trait-mediated indirect interactions: Primacy of trait-mediated indirect interactions. Ecology Letters 7:153–163.

Short, K. H., and K. Petren. 2008. Boldness underlies foraging success of invasive Lepidodactylus lugubris geckos in the human landscape. Animal Behaviour 76:429–437.

Siefert, A., C. Violle, L. Chalmandrier, C. H. Albert, A. Taudiere, A. Fajardo, L. W. Aarssen, et al. 2015. A global meta-analysis of the relative extent of intraspecific trait variation in plant communities. Ecology Letters 18:1406–1419.

Sih, A., A. Bell, and J. C. Johnson. 2004. Behavioral syndromes: an ecological and evolutionary overview. Trends in Ecology & Evolution 19:372–378.

Sobral, M. 2021. All Traits Are Functional: An Evolutionary Viewpoint. Trends in Plant Science 26:674–676.

Stamps, J. A., M. Briffa, and P. A. Biro. 2012. Unpredictable animals: individual differences in intraindividual variability (IIV). Animal Behaviour 83:1325–1334.

Szangolies, L., F. Jeltsch, L. G. Halsey, and C. A. Gallagher. 2025. From metabolism to coexistence: Understanding animal movement and community dynamics through energy. Individual-based Ecology 1:e151741.

Thompson, D. J., and O. M. Fincke. 2002. Body size and fitness in Odonata, stabilising selection and a meta-analysis too far? Ecological Entomology 27:378–384.

Toscano, B. J., N. J. Gownaris, S. M. Heerhartz, and C. J. Monaco. 2016. Personality, foraging behavior and specialization: integrating behavioral and food web ecology at the individual level. Oecologia 182:55–69.

Toscano, B. J., and B. D. Griffen. 2014. Trait-mediated functional responses: predator behavioural type mediates prey consumption. Journal of Animal Ecology 83:1469–1477.

Van Valen, L. 1965. Morphological Variation and Width of Ecological Niche. The American Naturalist 99:377–390.

Vanni, M. J., and P. B. McIntyre. 2016. Predicting nutrient excretion of aquatic animals with metabolic ecology and ecological stoichiometry: a global synthesis. Ecology 97:3460–3471.

Viechtbauer, W. 2010. Conducting Meta-Analyses in *R* with the **metafor** Package. Journal of Statistical Software 36.

Violle, C., B. J. Enquist, B. J. McGill, L. Jiang, C. H. Albert, C. Hulshof, V. Jung, et al. 2012. The return of the variance: intraspecific variability in community ecology. Trends in Ecology & Evolution 27:244–252.

Violle, C., M.-L. Navas, D. Vile, E. Kazakou, C. Fortunel, I. Hummel, and E. Garnier. 2007. Let the concept of trait be functional! Oikos 116:882–892.

Wainwright, P. C., and S. M. Reilly, eds. 1994. Ecological morphology: integrative organismal biology. University of Chicago Press, Chicago.

Wheelwright, N. T., L. F. Keller, and E. Postma. 2014. The effect of trait type and strength of selection on heritability and evolvability in an island bird population. Evolution 68:3325–3336.

Wong, M. K. L., and C. P. Carmona. 2021. Including intraspecific trait variability to avoid distortion of functional diversity and ecological inference: Lessons from natural assemblages. Methods in Ecology and Evolution. 12: 946–957.

Wray, M. K., H. R. Mattila, and T. D. Seeley. 2011. Collective personalities in honeybee colonies are linked to colony fitness. Animal Behaviour 81:559–568.

Wright, I. J., P. B. Reich, M. Westoby, D. D. Ackerly, Z. Baruch, F. Bongers, J. Cavender-Bares, et al. 2004. The worldwide leaf economics spectrum. Nature 428:821–827.

Wright, J., G. H. Bolstad, Y. G. Araya-Ajoy, and N. J. Dingemanse. 2019. Life-history evolution under fluctuating density-dependent selection and the adaptive alignment of pace-of-life syndromes: Pace-of-life syndromes. Biological Reviews 94:230–247.

Yang, Y., M. Lagisz, C. Williams, D. W. A. Noble, J. Pan, and S. Nakagawa. 2024. Robust point and variance estimation for meta-analyses with selective reporting and dependent effect sizes. Methods in Ecology and Evolution 15:1593–1610.

Zhao, T., S. Villéger, S. Lek, and J. Cucherousset. 2014. High intraspecific variability in the functional niche of a predator is associated with ontogenetic shift and individual specialization. Ecology and Evolution 4:4649–4657.

## Literature for the Meta-Analysis

Adams, C. E., and F. A. Huntingford. 2002. The functional significance of inherited differences in feeding morphology in a sympatric polymorphic population of Arctic charr. Evolutionary Ecology 16:15–25.

Akçay, Ç., S. E. Campbell, and M. D. Beecher. 2015. The fitness consequences of honesty: under-signalers have a survival advantage in song sparrows. Evolution 69:3186–3193.

Allgeier, J. E., T. J. Cline, T. E. Walsworth, G. Wathen, C. A. Layman, and D. E. Schindler. 2020. Individual behavior drives ecosystem function and the impacts of harvest. Science Advances 6:eaax8329.

Álvarez, D., and A. G. Nicieza. 2005. Is metabolic rate a reliable predictor of growth and survival of brown trout (Salmo trutta) in the wild? Canadian Journal of Fisheries and Aquatic Sciences 62:643–649.

Angelier, F., J. C. Wingfield, H. Weimerskirch, and O. Chastel. 2010. Hormonal correlates of individual quality in a long-lived bird: a test of the ‘corticosterone–fitness hypothesis’. Biology Letters 6:846–849.

Armitage, K. B. 1986. Individuality, social behavior, and reproductive success in yellow-bellied marmots. Ecology 67:1186–1193.

Armitage, K. B., and D. H. Van Vuren. 2003. Individual differences and reproductive success in yellow-bellied marmots. Ethology Ecology & Evolution 15:207–233.

Auer, S. K., K. Salin, G. J. Anderson, and N. B. Metcalfe. 2015a. Aerobic scope explains individual variation in feeding capacity. Biology Letters 11:20150793.

Auer, S. K., K. Salin, A. M. Rudolf, G. J. Anderson, and N. B. Metcalfe. 2015b. Flexibility in metabolic rate confers a growth advantage under changing food availability. Journal of Animal Ecology 84:1405–1411.

Baker, J. A., W. A. Cresko, S. A. Foster, and D. C. Heins. 2005. Life-history differentiation of benthic and limnetic ecotypes in a polytypic population of threespine stickleback (Gasterosteus aculeatus). Evolutionary Ecology Research 7:121–131.

Banks, P. B., K. Norrdahl, and E. Korpimäki. 2002. Mobility decisions and the predation risks of reintroduction. Biological Conservation 103:133–138.

Beauchamp, G. 2006. Phenotypic correlates of scrounging behavior in zebra finches: role of foraging efficiency and dominance. Ethology 112:873–878.

Berchtold, A. E., S. F. Colborne, F. J. Longstaffe, and B. D. Neff. 2015. Ecomorphological patterns linking morphology and diet across three populations of pumpkinseed sunfish (Lepomis gibbosus). Canadian Journal of Zoology 93:289–297.

Blackmer, A. L., R. A. Mauck, J. T. Ackerman, C. E. Huntington, G. A. Nevitt, and J. B. Williams. 2005. Exploring individual quality: basal metabolic rate and reproductive performance in storm-petrels. Behavioral Ecology 16:906–913.

Blake, C. A., and C. R. Gabor. 2014. Effect of prey personality depends on predator species. Behavioral Ecology 25:871–877.

Blanckenhorn, W. U. 1991a. Fitness consequences of food-based territoriality in water striders, Gerris remigis. Animal Behaviour 42:147–149.

Blanckenhorn, W. V. 1991b. Fitness consequences of foraging success in water striders (Gerris remigis; Heteroptera: Gerridae). Behavioral Ecology 2:46–55.

Blight, O., G. A. D. Mariblanca, X. Cerdá, and R. Boulay. 2016. A proactive–reactive syndrome affects group success in an ant species. Behavioral Ecology 27:118–125.

Bolnick, D. I., and M. S. Araújo. 2011. Partitioning the relative fitness effects of diet and trophic morphology in the threespine stickleback. Evolutionary Ecology Research 13:439–459.

Bolnick, D. I., and J. S. Paull. 2009. Morphological and dietary differences between individuals are weakly but positively correlated within a population of threespine stickleback. Evolutionary Ecology Research 11:1217–1233.

Boon, A. K., D. Réale, and S. Boutin. 2008. Personality, habitat use, and their consequences for survival in North American red squirrels Tamiasciurus hudsonicus. Oikos 117:1321–1328.

Boratyński, Z., E. Koskela, T. Mappes, and E. Schroderus. 2013. Quantitative genetics and fitness effects of basal metabolism. Evolutionary Ecology 27:301–314.

Boratyński, Z., E. Koskela, T. Mappes, and T. A. Oksanen. 2010. Sex-specific selection on energy metabolism—selection coefficients for winter survival. Journal of Evolutionary Biology 23:1969–1978.

Both, C., N. J. Dingemanse, P. J. Drent, and J. M. Tinbergen. 2005. Pairs of extreme avian personalities have highest reproductive success. Journal of Animal Ecology 74:667–674.

Boulton, K., C. A. Walling, A. J. Grimmer, G. G. Rosenthal, and A. J. Wilson. 2018. Phenotypic and genetic integration of personality and growth under competition in the sheepshead swordtail, Xiphophorus birchmanni. Evolution 72:187–201.

Bremner-Harrison, S., P. A. Prodohl, and R. W. Elwood. 2004. Behavioural trait assessment as a release criterion: boldness predicts early death in a reintroduction programme of captive-bred swift fox (Vulpes velox). Animal Conservation 7:313–320.

Brodin, T., and F. Johansson. 2004. Conflicting selection pressures on the growth/predation-risk trade-off in a damselfly. Ecology 85:2927–2932.

Calsbeek, R., C. Bonneaud, and T. B. Smith. 2008. Differential fitness effects of immunocompetence and neighbourhood density in alternative female lizard morphs. Journal of Animal Ecology 77:103–109.

Carlstead, K., J. Mellen, and D. G. Kleiman. 1999. Black rhinoceros (Diceros bicornis) in U.S. zoos: I. Individual behavior profiles and their relationship to breeding success. Zoo Biology 18:17–34.

Carter, A. J., A. W. Goldizen, and S. A. Tromp. 2010. Agamas exhibit behavioral syndromes: bolder males bask and feed more but may suffer higher predation. Behavioral Ecology 21:655–661.

Chism, J., and W. Rogers. 2010. Male competition, mating success and female choice in a seasonally breeding primate (Erythrocebus patas). Ethology 103:109–126.

Clobert, J., A. Oppliger, G. Sorci, B. Ernande, J. G. Swallow, and T. Garland. 2000. Trade-offs in phenotypic traits: endurance at birth, growth, survival, predation and susceptibility to parasitism in a lizard, Lacerta vivipara. Functional Ecology 14:675–684.

Cole, E. F., and J. L. Quinn. 2012. Personality and problem-solving performance explain competitive ability in the wild. Proceedings of the Royal Society B: Biological Sciences 279:1168–1175.

Cornwell, T. O., I. D. McCarthy, and P. A. Biro. 2020. Integration of physiology, behaviour and life history traits: personality and pace of life in a marine gastropod. Animal Behaviour 163:155–162.

Cote, J., A. Dreiss, and J. Clobert. 2008. Social personality trait and fitness. Proceedings of the Royal Society B: Biological Sciences 275:2851–2858.

Cucherousset, J., A. Acou, S. Blanchet, J. R. Britton, W. R. C. Beaumont, and R. E. Gozlan. 2011. Fitness consequences of individual specialisation in resource use and trophic morphology in European eels. Oecologia 167:75–84.

Cutts, C. J., C. E. Adams, and A. Campbell. 2001. Stability of physiological and behavioural determinants of performance in Arctic char (Salvelinus alpinus). Canadian Journal of Fisheries and Aquatic Sciences 58:961–968.

David, M., Y. Auclair, and F. Cézilly. 2011. Personality predicts social dominance in female zebra finches, Taeniopygia guttata, in a feeding context. Animal Behaviour 81:219–224.

David, M., Y. Auclair, L.-A. Giraldeau, and F. Cézilly. 2012. Personality and body condition have additive effects on motivation to feed in zebra finches Taeniopygia guttata. Ibis 154:372–378.

Day, T., and J. D. McPhail. 1996. The effect of behavioural and morphological plasticity on foraging efficiency in the threespine stickleback (Gasterosteus sp.). Oecologia 108:380–388.

Denoël, M. 2004. Feeding performance in heterochronic alpine newts is consistent with trophic niche and maintenance of polymorphism. Ethology 110:127–136.

Derting, T. L., and P. A. McClure. 1989. Intraspecific variation in metabolic rate and its relationship with productivity in the cotton rat, Sigmodon hispidus. Journal of Mammalogy 70:520–531.

Des Roches, S., J. B. Shurin, D. Schluter, and L. J. Harmon. 2013. Ecological and evolutionary effects of stickleback on community structure. PLoS ONE 8:e59644.

Dewsbury, D. A. 1984. Aggression, copulation, and differential reproduction of deer mice (Peromyscus maniculatus) in a semi-natural enclosure. Behaviour 91:1–23.

Diaz Pauli, B., E. Edeline, and C. Evangelista. 2020. Ecosystem consequences of multi-trait response to environmental changes in Japanese medaka, Oryzias latipes. Conservation Physiology 8:coaa011.

Dingemanse, N. J., C. Both, P. J. Drent, and J. M. Tinbergen. 2004. Fitness consequences of avian personalities in a fluctuating environment. Proceedings of the Royal Society of London B: Biological Sciences 271:847–852.

Dugatkin, L. A. 1992. Tendency to inspect predators predicts mortality risk in the guppy (Poecilia reticulata). Behavioral Ecology 3:124–127.

Ehlinger, T. J., and D. S. Wilson. 1988. Complex foraging polymorphism in bluegill sunfish. Proceedings of the National Academy of Sciences 85:1878–1882.

Einum, S., E. I. F. Fossen, V. Parry, and C. Pélabon. 2019. Genetic variation in metabolic rate and correlations with other energy budget components and life history in Daphnia magna. Evolutionary Biology 46:170–178.

Fernández-Reiriz, M. J., J. Irisarri, and U. Labarta. 2016. Flexibility of physiological traits underlying inter-individual growth differences in intertidal and subtidal mussels Mytilus galloprovincialis. PLoS ONE 11:e0148245.

Fisher, D. N., M. David, R. Rodríguez-Muñoz, and T. Tregenza. 2018. Lifespan and age, but not residual reproductive value or condition, are related to behaviour in wild field crickets. Ethology 282:338–346.

Foote, A. D., J. Newton, S. B. Piertney, E. Willerslev, and M. T. P. Gilbert. 2009. Ecological, morphological and genetic divergence of sympatric North Atlantic killer whale populations. Molecular Ecology 18:5207–5217.

Formica, V. A., C. W. Wood, W. B. Larsen, R. E. Butterfield, M. E. Augat, H. Y. Hougen, and E. D. Brodie. 2012. Fitness consequences of social network position in a wild population of forked fungus beetles (Bolitotherus cornutus). Journal of Evolutionary Biology 25:130–137.

Foster, W. C., C. M. Armstrong, G. T. Chism, and J. N. Pruitt. 2017. Smaller and bolder prey snails have higher survival in staged encounters with the sea star Pisaster giganteus. Current Zoology, zow116.

Gagnon, M.-C., and J. Turgeon. 2011. Sexual conflict in Gerris gillettei (Insecta: Hemiptera): intraspecific intersexual correlated morphology and experimental assessment of behaviour and fitness. Journal of Evolutionary Biology 24:1505–1516.

Galeotti, P., R. Sacchi, D. Pellitteri-Rosa, A. Bellati, W. Cocca, A. Gentilli, S. Scali, and M. Fasola. 2013. Colour polymorphism and alternative breeding strategies: effects of parent’s colour morph on fitness traits in the common wall lizard. Evolutionary Biology 40:385–394.

Gangloff, E. J., D. Vleck, and A. M. Bronikowski. 2015. Developmental and immediate thermal environments shape energetic trade-offs, growth efficiency, and metabolic rate in divergent life-history ecotypes of the garter snake Thamnophis elegans. Physiological and Biochemical Zoology 88:550–563.

Gifford, M. E., T. A. Clay, and V. Careau. 2014. Individual (co)variation in standard metabolic rate, feeding rate, and exploratory behavior in wild-caught semiaquatic salamanders. Physiological and Biochemical Zoology 87:384–396.

Giménez, J. O. 2015. Determinants of phenotypic variation in the Iberian wall lizard species complex (Podarcis hispanicus). Master’s thesis.

Glon, M. G., E. R. Larson, and K. L. Pangle. 2016. Connecting laboratory behavior to field function through stable isotope analysis. PeerJ 4:e1918.

Griffen, B. D., and H. Mosblack. 2011. Predicting diet and consumption rate differences between and within species using gut ecomorphology. Journal of Animal Ecology 80:854–863.

Hromada, M., L. Kuczyński, A. Krištín, and P. Tryjanowski. 2003. Animals of different phenotype differentially utilise dietary niche—the case of the great grey shrike Lanius excubitor. Ornis Fennica 80:71–78.

Hsu, Y.-C., P.-J. Shaner, C.-I. Chang, L. Ke, and S.-J. Kao. 2014. Trophic niche width increases with bill-size variation in a generalist passerine: a test of niche variation hypothesis. Journal of Animal Ecology 83:450–459.

Hulthén, K., B. B. Chapman, P. A. Nilsson, L.-A. Hansson, C. Skov, J. Brodersen, J. Vinterstare, and C. Brönmark. 2017. A predation cost to bold fish in the wild. Scientific Reports 7:1239.

Ingram, T., R. Svanbäck, N. J. B. Kraft, P. Kratina, L. Southcott, and D. Schluter. 2012. Intraguild predation drives evolutionary niche shift in threespine stickleback. Evolution 66:1819–1832.

Ioannou, C. C., M. Payne, and J. Krause. 2008. Ecological consequences of the bold–shy continuum: the effect of predator boldness on prey risk. Oecologia 157:177.

Jablonszky, M., E. Szász, K. Krenhardt, G. Markó, G. Hegyi, M. Herényi, M. Laczi, G. Nagy, B. Rosivall, E. Szöllősi, J. Török, and L. Z. Garamszegi. 2018. Unravelling the relationships between life history, behaviour and condition under the pace-of-life syndromes hypothesis using long-term data from a wild bird. Behavioral Ecology and Sociobiology 72:52.

Johnson, J. C., and A. Sih. 2005. Precopulatory sexual cannibalism in fishing spiders (Dolomedes triton): a role for behavioral syndromes. Behavioral Ecology and Sociobiology 58:390–396.

Jolles, J. W., A. Manica, and N. J. Boogert. 2016. Food intake rates of inactive fish are positively linked to boldness in three-spined sticklebacks Gasterosteus aculeatus. Journal of Fish Biology 88:1661–1668.

Kain, M. P., and M. W. McCoy. 2016. Anti-predator behavioral variation among Physa acuta in response to temporally fluctuating predation risk by Procambarus. Behavioural Processes 133:15–23.

Karpestam, E., and A. Forsman. 2011. Dietary differences among colour morphs of pygmy grasshoppers revealed by behavioural experiments and stable isotopes. Evolutionary Ecology Research 13:461–477.

Katano, O. 2011. Effects of individual differences in foraging of pale chub on algal biomass through trophic cascades. Environmental Biology of Fishes 92:101–112.

Kause, A., I. Saloniemi, E. Haukioja, and S. Hanhimäki. 1999. How to become large quickly: quantitative genetics of growth and foraging in a flush feeding lepidopteran larva. Journal of Evolutionary Biology 12:471–482.

Keiser, C. N., S. J. Ingley, B. J. Toscano, I. Scharf, and J. N. Pruitt. 2018. Habitat complexity dampens selection on prey activity level. Ethology 124:25–32.

Keiser, C. N., J. B. Slyder, W. P. Carson, and J. N. Pruitt. 2015. Individual differences in predators but not producers mediate the magnitude of a trophic cascade. Arthropod-Plant Interactions 9:225–232.

Kerr, N. R. 2017. Links between personality and individual niche in the freshwater fish Gobiomorphus cotidianus. Master’s thesis, Otago, New Zealand.

Khelifa, R., R. Zebsa, H. Amari, M. K. Mellal, and H. Mahdjoub. 2019. Field estimates of fitness costs of the pace-of-life in an endangered damselfly. Journal of Evolutionary Biology 32:943–954.

Kobler, A., T. Klefoth, T. Mehner, and R. Arlinghaus. 2009. Coexistence of behavioural types in an aquatic top predator: a response to resource limitation? Oecologia 161:837–847.

Korhonen, H., P. Niemelä, and P. Siirilä. 2001. Temperament and reproductive performance in farmed sable. Agricultural Food Science Finland 10:91–98.

Korhonen, H. T., L. Jauhiainen, and T. Rekilä. 2002. Effect of temperament and behavioural reactions to the presence of a human during the pre-mating period on reproductive performance in farmed mink (Mustela vison). Canadian Journal of Animal Science 82:275–282.

Kralj-Fišer, S., E. A. Hebets, and M. Kuntner. 2017. Different patterns of behavioral variation across and within species of spiders with differing degrees of urbanization. Behavioral Ecology and Sociobiology 71:125.

Kruuk, L. E. B., T. H. Clutton-Brock, J. Slate, J. M. Pemberton, S. Brotherstone, and F. E. Guinness. 2000. Heritability of fitness in a wild mammal population. Proceedings of the National Academy of Sciences 97:698–703.

Lapiedra, O., T. W. Schoener, M. Leal, J. B. Losos, and J. J. Kolbe. 2018. Predator-driven natural selection on risk-taking behavior in anole lizards. Science 360:1017–1020.

Larivée, M., S. Boutin, J. R. Speakman, and A. G. McAdam. 2010. Associations between over-winter survival and resting metabolic rate in juvenile North American red squirrels. Functional Ecology 27:597–607.

Lattanzio, M. S., and D. B. Miles. 2016. Trophic niche divergence among colour morphs that exhibit alternative mating tactics. Royal Society Open Science 3:150531.

Laurila, A., and J. Kujasalo. 1999. Habitat duration, predation risk and phenotypic plasticity in common frog (Rana temporaria) tadpoles. Journal of Animal Ecology 68:1123–1132.

Lavin, P. A., and J. D. McPhail. 1986. Adaptive divergence of trophic phenotype among freshwater populations of the threespine stickleback (Gasterosteus aculeatus). Canadian Journal of Fisheries and Aquatic Sciences 43:2455–2463.

Lepetz, V., M. Massot, A. S. Chaine, and J. Clobert. 2009. Climate warming and the evolution of morphotypes in a reptile. Global Change Biology 15:454–466.

Luiz, O. J., D. A. Crook, M. J. Kennard, J. D. Olden, T. M. Saunders, M. M. Douglas, D. Wedd, and A. J. King. 2019. Does a bigger mouth make you fatter? Linking intraspecific gape variability to body condition of a tropical predatory fish. Oecologia 191:579–585.

Maldonado, K., S. Pablo, G. Piriz, J. M. Bogdanovich, R. F. Nespolo, and F. Bozinovic. 2016. Is maximum food intake in endotherms constrained by net or factorial aerobic scope? Lessons from the leaf-eared mouse. Frontiers in Physiology 7.

Marshall, H. H., J. L. Sanderson, F. Mwanghuya, R. Businge, S. Kyabulima, M. C. Hares, E. Inzani, G. Kalema-Zikusoka, K. Mwesige, F. J. Thompson, E. I. K. Vitikainen, and M. A. Cant. 2016. Variable ecological conditions promote male helping by changing banded mongoose group composition. Behavioral Ecology 27:978–987.

Maskrey, D. K., S. J. White, A. J. Wilson, and T. M. Houslay. 2018. Who dares does not always win: risk-averse rockpool prawns are better at controlling a limited food resource. Animal Behaviour 140:187–197.

Matthews, B., T. Aebischer, K. E. Sullam, B. Lundsgaard-Hansen, and O. Seehausen. 2016. Experimental evidence of an eco-evolutionary feedback during adaptive divergence. Current Biology 26:483–489.

Matthews, B., K. B. Marchinko, D. I. Bolnick, and A. Mazumder. 2010. Specialization of trophic position and habitat use by sticklebacks in an adaptive radiation. Ecology 91:1025–1034.

McCollum, S. A., and J. Van Buskirk. 1996. Costs and benefits of a predator-induced polyphenism in the gray treefrog Hyla chrysoscelis. Evolution 50:583–593.

McPhee, M. V., and T. P. Quinn. 1998. Factors affecting the duration of nest defense and reproductive lifespan of female sockeye salmon, Oncorhynchus nerka. Environmental Biology of Fishes 51:369–375.

Metcalfe, N. B., A. C. Taylor, and J. E. Thorpe. 1995. Metabolic rate, social status and life-history strategies in Atlantic salmon. Animal Behaviour 49:431–436.

Møller, A. P. 2010. Brain size, head size and behaviour of a passerine bird. Journal of Evolutionary Biology 23:625–635.

Monceau, K., F.-X. Dechaume-Moncharmont, J. Moreau, C. Lucas, R. Capoduro, S. Motreuil, and Y. Moret. 2017. Personality, immune response and reproductive success: an appraisal of the pace-of-life syndrome hypothesis. Journal of Animal Ecology 86:932–942.

Montiglio, P., T. W. Wey, A. T. Chang, S. Fogarty, and A. Sih. 2017. Correlational selection on personality and social plasticity: morphology and social context determine behavioural effects on mating success. Journal of Animal Ecology 86:213–226.

Morales, J. A., D. G. Cardoso, T. M. C. Della Lucia, and R. N. C. Guedes. 2013. Weevil x insecticide: does ‘personality’ matter? PLoS ONE 8:e67283.

Nakayama, S., and L. A. Fuiman. 2010. Body size and vigilance mediate asymmetric interference competition for food in fish larvae. Behavioral Ecology 21:708–713.

Nakayama, S., T. Rapp, and R. Arlinghaus. 2017. Fast–slow life history is correlated with individual differences in movements and prey selection in an aquatic predator in the wild. Journal of Animal Ecology 86:192–201.

Nannini, M. A., J. Parkos, and D. H. Wahl. 2012. Do behavioral syndromes affect foraging strategy and risk-taking in a juvenile fish predator? Transactions of the American Fisheries Society 141:26–33.

Niemelä, P. T., E. Z. Lattenkamp, and N. J. Dingemanse. 2015. Personality-related survival and sampling bias in wild cricket nymphs. Behavioral Ecology 26:936–946.

Niemelä, P. T., P. P. Niehoff, C. Gasparini, N. J. Dingemanse, and C. Tuni. 2019. Crickets become behaviourally more stable when raised under higher temperatures. Behavioral Ecology and Sociobiology 73:81.

Noble, D. W. A., K. Wechmann, J. S. Keogh, and M. J. Whiting. 2013. Behavioral and morphological traits interact to promote the evolution of alternative reproductive tactics in a lizard. American Naturalist 182:726–742.

Nyqvist, M. J., J. Cucherousset, R. E. Gozlan, and J. R. Britton. 2018. Relationships between individual movement, trophic position and growth of juvenile pike (Esox lucius). Ecology of Freshwater Fish 27:398–407.

Oretga, J., D. Pellitteri-Rosa, P. López, and J. Martín. 2015. Dorsal pattern polymorphism in female Iberian wall lizards: differences in morphology, dorsal coloration, immune response, and reproductive investment. Biological Journal of the Linnean Society 116:352–363.

Ouyang, J. Q., P. J. Sharp, A. Dawson, M. Quetting, and M. Hau. 2011. Hormone levels predict individual differences in reproductive success in a passerine bird. Proceedings of the Royal Society B: Biological Sciences 278:2537–2545.

Palkovacs, E. P., and D. M. Post. 2009. Experimental evidence that phenotypic divergence in predators drives community divergence in prey. Ecology 90:300–305.

Palkovacs, E. P., B. A. Wasserman, and M. T. Kinnison. 2011. Eco-evolutionary trophic dynamics: loss of top predators drives trophic evolution and ecology of prey. PLoS ONE 6:e18879.

Parsons, K. J., and B. W. Robinson. 2007. Foraging performance of diet-induced morphotypes in pumpkinseed sunfish (Lepomis gibbosus) favours resource polymorphism. Journal of Evolutionary Biology 20:673–684.

Patrick, S. C., and H. Weimerskirch. 2014. Personality, foraging and fitness consequences in a long lived seabird. PLoS ONE 9:e87269.

Peiman, K. S., K. Birnie-Gauvin, M. H. Larsen, S. F. Colborne, K. M. Gilmour, K. Aarestrup, W. G. Willmore, and S. J. Cooke. 2017. Morphological, physiological and dietary covariation in migratory and resident adult brown trout (Salmo trutta). Zoology 123:79–90.

Petitjean, Q., S. Jean, J. Côte, A. Lamarins, M. Lefranc, R. Santos, A. Perrault, P. Laffaille, and L. Jacquin. 2020. Combined effects of temperature increase and immune challenge in two wild gudgeon populations. Fish Physiology and Biochemistry 46:157–176.

Pintor, L. M., K. E. McGhee, D. P. Roche, and A. M. Bell. 2014. Individual variation in foraging behavior reveals a trade-off between flexibility and performance of a top predator. Behavioral Ecology and Sociobiology 68:1711–1722.

Pintor, L. M., A. Sih, and M. L. Bauer. 2008. Differences in aggression, activity and boldness between native and introduced populations of an invasive crayfish. Oikos 117:1629–1636.

Pintor, L. M., A. Sih, and J. L. Kerby. 2009. Behavioral correlations provide a mechanism for explaining high invader densities and increased impacts on native prey. Ecology 90:581–587.

Piquet, J. C., M. López-Darias, A. van der Marel, M. Nogales, and J. Waterman. 2018. Unraveling behavioral and pace-of-life syndromes in a reduced parasite and predation pressure context: personality and survival of the Barbary ground squirrel. Behavioral Ecology and Sociobiology 72:147.

Powers, D. R., and K. A. Nagy. 1988. Field metabolic rate and food consumption by free-living Anna’s hummingbirds (Calypte anna). Physiological Zoology 61:500–506.

Pruitt, J. N., C. N. Keiser, B. Banka, J. S. Liedle, A. J. Brooks, R. J. Schmitt, and S. J. Holbrook. 2018. Collective aggressiveness of an ecosystem engineer is associated with coral recovery. Behavioral Ecology.

Pruitt, J. N., S. E. Riechert, and T. C. Jones. 2008. Behavioural syndromes and their fitness consequences in a socially polymorphic spider, Anelosimus studiosus. Animal Behaviour 76:871–879.

Quevedo, M., R. Svanbäck, and P. Eklöv. 2009. Intrapopulation niche partitioning in a generalist predator limits food web connectivity. Ecology 90:2263–2274.

Quinn, J. L., S. C. Patrick, S. Bouwhuis, T. A. Wilkin, and B. C. Sheldon. 2009. Heterogeneous selection on a heritable temperament trait in a variable environment. Journal of Animal Ecology 78:1203–1215.

Raffard, A., J. Cucherousset, F. Santoul, L. Di Gesu, and S. Blanchet. 2023. Climate and intraspecific variation in a consumer species drive ecosystem multifunctionality. Oikos. e09286.

Raffard, A., F. Santoul, S. Blanchet, and J. Cucherousset. 2020. Linking intraspecific variability in trophic and functional niches along an environmental gradient. Freshwater Biology 65:1401–1411.

Raine, N. E., and L. Chittka. 2005. Colour preferences in relation to the foraging performance and fitness of the bumblebee Bombus terrestris. Uludag Bee Journal 5:145–150.

Réale, D., and M. Festa-Bianchet. 2003. Predator-induced natural selection on temperament in bighorn ewes. Animal Behaviour 65:463–470.

Réale, D., B. Y. Gallant, M. Leblanc, and M. Festa-Bianchet. 2000. Consistency of temperament in bighorn ewes and correlates with behaviour and life history. Animal Behaviour 60:589–597.

Reid, D., J. D. Armstrong, and N. B. Metcalfe. 2011. Estimated standard metabolic rate interacts with territory quality and density to determine the growth rates of juvenile Atlantic salmon. Functional Ecology 25:1360–1367.

Reid, D., J. D. Armstrong, and N. B. Metcalfe. 2012. The performance advantage of a high resting metabolic rate in juvenile salmon is habitat dependent. Journal of Animal Ecology 81:868–875.

Robinson, B. W. 2000. Trade offs in habitat-specific foraging efficiency and the nascent adaptive divergence of sticklebacks in lakes. Behaviour 137:865–888.

Rota, T., J. Jabiol, S. Lamothe, D. Lambrigot, É. Chauvet, and A. Lecerf. 2024. Phenotypes of predator individuals underpin contrasting ecosystem effects. Freshwater Biology 69:739–753.

Rudman, S. M., M. A. Rodriguez-Cabal, A. Stier, T. Sato, J. Heavyside, R. W. El-Sabaawi, and G. M. Crutsinger. 2015. Adaptive genetic variation mediates bottom-up and top-down control in an aquatic ecosystem. Proceedings of the Royal Society B: Biological Sciences 282:20151234.

Rudman, S. M., and D. Schluter. 2016. Ecological impacts of reverse speciation in threespine stickleback. Current Biology 26:490–495.

Sampaio, A. L. A., J. P. A. Pagotto, and E. Goulart. 2013. Relationships between morphology, diet and spatial distribution: testing the effects of intra and interspecific morphological variations on the patterns of resource use in two Neotropical cichlids. Neotropical Ichthyology 11:351–360.

Santicchia, F., C. Gagnaison, F. Bisi, A. Martinoli, E. Matthysen, S. Bertolino, and L. A. Wauters. 2018. Habitat-dependent effects of personality on survival and reproduction in red squirrels. Behavioral Ecology and Sociobiology 72:134.

Santostefano, F., A. J. Wilson, P. T. Niemelä, and N. J. Dingemanse. 2017. Behavioural mediators of genetic life-history trade-offs: a test of the pace-of-life syndrome hypothesis in field crickets. Proceedings of the Royal Society B: Biological Sciences 284:20171567.

Sarno, R. J., and W. L. Franklin. 1999. Maternal expenditure in the polygynous and monomorphic guanaco: suckling behavior, reproductive effort, yearly variation, and influence on juvenile survival. Behavioral Ecology 10:41–47.

Schaack, S., and L. J. Chapman. 2003. Interdemic variation in the African cyprinid Barbus neumayeri: correlations among hypoxia, morphology, and feeding performance. Canadian Journal of Zoology 81:430–440.

Schuett, W., S. R. X. Dall, M. H. Kloesener, J. Baeumer, F. Beinlich, and T. Eggers. 2015. Life-history trade-offs mediate ‘personality’ variation in two colour morphs of the pea aphid, Acyrthosiphon pisum. Journal of Animal Ecology 84:90–101.

Schulte-Hostedde, A. I., and J. S. Millar. 2004. Intraspecific variation of testis size and sperm length in the yellow-pine chipmunk (Tamias amoenus): implications for sperm competition and reproductive success. Behavioral Ecology and Sociobiology 55:272–277.

Shackleton, M. A., M. D. Jennions, and J. Hunt. 2005. Fighting success and attractiveness as predictors of male mating success in the black field cricket, Teleogryllus commodus: the effectiveness of no-choice tests. Behavioral Ecology and Sociobiology 58:1–8.

Smith, T. B. 1987. Bill size polymorphism and intraspecific niche utilization in an African finch. Nature 329:717–719.

Snowberg, L. K., and D. I. Bolnick. 2008. Assortative mating by diet in a phenotypically unimodal but ecologically variable population of stickleback. American Naturalist 172:733–739.

Snowberg, L. K., K. M. Hendrix, and D. I. Bolnick. 2015. Covarying variances: more morphologically variable populations also exhibit more diet variation. Oecologia 178:89–101.

Soares, A. O., and D. C. H. Schanderl. 2001. Fitness of two phenotypes of Harmonia axyridis (Coleoptera: Coccinellidae). 98:287–293.

Song, Z.-G., and D.-H. Wang. 2002. The maximum metabolizable energy intake and the relationship with basal metabolic rate in the striped hamster Cricetulus barabensis. Acta Theriologica 47:417–423.

Speakman, J. R., E. Król, and M. S. Johnson. 2004. The functional significance of individual variation in basal metabolic rate. Physiological and Biochemical Zoology 77:900–915.

Spritzer, M. D., D. B. Meikle, and N. G. Solomon. 2005. Female choice based on male spatial ability and aggressiveness among meadow voles. Animal Behaviour 69:1121–1130.

Start, D., and B. Gilbert. 2017. Predator personality structures prey communities and trophic cascades. Ecology Letters 20:366–374.

Start, D., and B. Gilbert. 2019. Trait variation across biological scales shapes community structure and ecosystem function. Ecology 100.

Steinmeyer, C., J. C. Mueller, and B. Kempenaers. 2013. Individual variation in sleep behaviour in blue tits Cyanistes caeruleus: assortative mating and associations with fitness-related traits. Journal of Avian Biology 44:159–168.

Svanbäck, R., and P. Eklöv. 2004. Morphology in perch affects habitat specific feeding efficiency. Functional Ecology 18:503–510.

Taylor, R. W., S. Boutin, M. M. Humphries, and A. G. McAdam. 2014. Selection on female behaviour fluctuates with offspring environment. Journal of Evolutionary Biology 27:2308–2321.

Van Buskirk, J., S. A. McCollum, and E. E. Werner. 1997. Natural selection for environmentally induced phenotypes in tadpoles. Evolution 51:1983–1992.

Van Buskirk, J., and S. A. McCollum. 2000. Functional mechanisms of an inducible defence in tadpoles: morphology and behaviour influence mortality risk from predation. Journal of Evolutionary Biology 13:336–347.

Van Overveld, T., F. Adriaensen, and E. Matthysen. 2015. No evidence for correlational selection on exploratory behaviour and natal dispersal in the great tit. Evolutionary Ecology 29:137–156.

Vargas, R., S. Mackenzie, and S. Rey. 2018. ’Love at first sight’: the effect of personality and colouration patterns in the reproductive success of zebrafish (Danio rerio). PLoS ONE 13:e0203320.

Vercken, E., M. Massot, B. Sinervo, and J. Clobert. 2007. Colour variation and alternative reproductive strategies in females of the common lizard Lacerta vivipara. Journal of Evolutionary Biology 20:221–232.

Walton, M. 1988. Relationships among metabolic, locomotory, and field measures of organismal performance in the Fowler’s toad (Bufo woodhousei fowleri). Physiological Zoology 61:107–118.

Ward, A. J. W., P. Thomas, P. J. B. Hart, and J. Krause. 2004. Correlates of boldness in three-spined sticklebacks (Gasterosteus aculeatus). Behavioral Ecology and Sociobiology 55:561–568.

Webster, M. M., A. J. W. Ward, and P. J. B. Hart. 2009. Individual boldness affects interspecific interactions in sticklebacks. Behavioral Ecology and Sociobiology 63:511–520.

Wellborn, G. A. 2000. Selection on a sexually dimorphic trait in ecotypes within the Hyalella azteca species complex (Amphipoda: Hyalellidae). American Midland Naturalist 143:212–225.

White, J. R., M. G. Meekan, M. I. McCormick, and M. C. O. Ferrari. 2013. A comparison of measures of boldness and their relationships to survival in young fish. PLoS ONE 8:e68900.

Whiteman, H. H., S. A. Wissinger, and W. S. Brown. 1996. Growth and foraging consequences of facultative paedomorphosis in the tiger salamander, Ambystoma tigrinum nebulosum. Evolutionary Ecology 10:433–446.

Wielebnowski, N. C. 1999. Behavioral differences as predictors of breeding status in captive cheetahs. Zoo Biology 18:335–349.

Wilson, A. D. M., J.-G. J. Godin, and A. J. W. Ward. 2010. Boldness and reproductive fitness correlates in the eastern mosquitofish, Gambusia holbrooki. Ethology 116:96–104.

Wilson, D. S., K. Coleman, A. B. Clark, and L. Biederman. 1993. Shy-bold continuum in pumpkinseed sunfish (Lepomis gibbosus): an ecological study of a psychological trait. Journal of Comparative Psychology 107:250–260.

Yamamoto, T., H. Ueda, and S. Higashi. 1998. Correlation among dominance status, metabolic rate and otolith size in masu salmon. Journal of Fish Biology 52:281–290.

Zavorka, L., D. Aldven, J. Naslund, J. Hojesjo, and J. I. Johnsson. 2015. Linking lab activity with growth and movement in the wild: explaining pace-of-life in a trout stream. Behavioral Ecology 26:877–884.

Zavorka, L., D. Aldven, J. Naslund, J. Hojesjo, and J. I. Johnsson. 2016. Inactive trout come out at night: behavioral variation, circadian activity, and fitness in the wild. Ecology 97:2223–2231.

Závorka, L., B. Koeck, J. Cucherousset, J. Brijs, J. Näslund, D. Aldvén, J. Höjesjö, I. A. Fleming, and J. I. Johnsson. 2017. Co-existence with non-native brook trout breaks down the integration of phenotypic traits in brown trout parr. Functional Ecology 31:1582–1591.

Zeng, L.-Q., A.-J. Zhang, S. S. Killen, Z.-D. Cao, Y.-X. Wang, and S.-J. Fu. 2017. Standard metabolic rate predicts growth trajectory of juvenile crucian carp (Carassius auratus) under changing food availability. Biology Open, bio.025452.

Zub, K., Z. Borowski, P. A. Szafrańska, M. Wieczorek, and M. Konarzewski. 2014. Lower body mass and higher metabolic rate enhance winter survival in root voles, Microtus oeconomus. Biological Journal of the Linnean Society 113:297–309.

Koricheva, J., J. Gurevitch, and K. Mengersen, eds. 2013. Handbook of meta-analysis in ecology and evolution. Princeton University Press, Princeton.

## References Cited Only in the Online Enhancements

Maino, J. L., and M. R. Kearney. 2015. Ontogenetic and interspecific scaling of consumption in insects. Oikos 124:1564–1570.

Ronget, V., J. Gaillard, T. Coulson, M. Garratt, F. Gueyffier, J. Lega, and J. Lemaître. 2018. Causes and consequences of variation in offspring body mass: meta-analyses in birds and mammals. Biological Reviews 93:1–27.

Rudolf, V. H. W., and N. L. Rasmussen. 2013. Population structure determines functional differences among species and ecosystem processes. Nature Communications 4:2318.

Smith, B. R., and D. T. Blumstein. 2008. Fitness consequences of personality: a meta-analysis. Behavioral Ecology 19:448–455.

