## Supplementary Materials for "Behavior and Physiology Outpace Form When Linking Traits to Ecological Responses Within Populations: A Meta-Analysis"

*Journal:* ***The American Naturalist***

***Section 1****: Enhanced informations*

**Table S1**: Detailed list of the 30 search term combinations (search strings) entered in each of the three search platforms (Web of Science ‘WoS’, Scopus and Google Scholar ‘GS’). The search started during year 2019 and ended up in 2020. We included studies until the end of our search in 2020. For GS searches, we delimited our searches within the first twenty pages (i.e., first 200 results based on relevance to our search terms).

| ***Search strings*** | ***WoS*** | ***Scopus*** | ***GS*** |  |
| --- | --- | --- | --- | --- |
| phenotypic  AND traits  AND individual  AND intraspecific  AND fitness  functional  AND traits  AND individual  AND intraspecific  AND fitness  morphological  AND traits  AND individual  AND intraspecific  AND fitness  physiological  AND traits  AND individual  AND intraspecific  AND fitness  behavioral AND traits  AND individual  AND intraspecific  AND fitness  phenotypic AND traits AND individual AND intraspecific AND feeding AND consumption rate  functional  AND traits AND individual AND intraspecific AND feeding AND consumption rate  morphological AND traits AND individual AND intraspecific AND feeding AND consumption rate  physiological AND traits AND individual AND intraspecific AND feeding AND consumption rate  behavioral AND traits AND individual AND intraspecific AND feeding AND consumption rate  phenotypic AND traits AND individual AND intraspecific AND community structure  functional  AND traits AND individual AND intraspecific AND community structure  morphological AND traits AND individual AND intraspecific AND community structure  physiological AND traits AND individual AND intraspecific AND community structure  behavioral AND traits AND individual AND intraspecific AND community structure  phenotypic AND traits AND individual AND intraspecific AND ecosystem functions  functional  AND traits AND individual AND intraspecific AND ecosystem functions  morphological AND traits AND individual AND intraspecific AND ecosystem functions  physiological AND traits AND individual AND intraspecific AND ecosystem functions  behavioral AND traits AND individual AND intraspecific AND ecosystem functions  phenotypic AND traits AND individual AND intraspecific AND isotopic niche  functional  AND traits AND individual AND intraspecific AND isotopic niche  morphological AND traits AND individual AND intraspecific AND isotopic niche  physiological AND traits AND individual AND intraspecific AND isotopic niche  behavioral AND traits AND individual AND intraspecific AND isotopic niche  phenotypic AND traits AND individual AND intraspecific AND trophic niche  functional  AND traits AND individual AND intraspecific AND trophic niche  morphological AND traits AND individual AND intraspecific AND trophic niche  physiological AND traits AND individual AND intraspecific AND trophic niche  behavioral AND traits AND individual AND intraspecific AND trophic niche | 76  38  13  77  24  1  1  2  1  0  37  73  11  12  8  26  54  13  6  6  1  1  1  1  0  10  11  4  1  2 | 53  33  15  29  36  2  0  1  1  2  25  44  8  10  8  19  46  13  11  8  1  1  1  2  1  6  8  3  3  3 | 200  200  200  200  200  200  200  200  200  200  200  200  200  200  200  200  200  200  200  200  200  200  200  200  200  200  200  200  200  200 |  |
| **TOTAL (with Duplicates)** | **511** | **393** | **6000** | **6904** |

**Box S1**. Decision path reporting how the potential dependence of effect sizes with body size or ontogeny was treated in each original study, alongside our associated confidence score (from zero to seven, with low values indicating poor confidence, and high values, high confidence).

A. In each primary study, we aimed that the relationship between the trait and the response was not resulting from variation in body size, so we included the effect size, if at least one of these two conditions was filled:

**A 1)** No strong relationship between body size and the response nor the trait was expected theoretically (e.g., by the Metabolic Theory of Ecology (Brown et al., 2004) or by life-history theory) (**A 1) alone = 2**).

**A 2)** The analyses were performed on a group of individuals or populations of a same size class, age or cohort (**A 2) alone = 2**).

B. For cases where body size was theoretically expected to affect both trait and response (e.g., body size links both to metabolic and feeding rate; Brown et al., 2004), we only extracted statistics :

(***i***) From partial relationships (i.e., body size has been accounted for on the response and the trait with regressions). (**B(*i*) alone = 3**).

(***ii***) When authors tested the relationships on mass-specific traits and/or mass-specific responses (i.e., in dividing the trait and/or the response by body size or in using allometric corrections). (**B(*ii*) alone = 2.5**).

C. When the effect of body size was expected on the response or on the trait only (e.g., body size was related to feeding rate, but not to boldness):

(***iii***) We also included statistics from semi-partial relationships (i.e., accounting for the dependent effect of body size on either the response or on the trait, for instance when authors added body size as a covariate in their model, but did not specify if the trait was size-corrected or not). (**C(*iii*) alone = 1.5**).

D. When trait-to-response relationships were obviously a result of variation in body size, or when we had a doubt, we did not include the effect size(s).
E. However, in rare cases, we kept studies that were part of previous meta-analyses which tested how behavior affected fitness (Smith and Blumstein 2008; Moiron et al. 2020), but for which the information on a potential dependence with body size was lacking (**confidence score = zero**). We attributed an average **confidence score of 4.5** for unpublished estimates we found in these two references.
F. When studies combined different methods to account for a potential bias due to the influence of body size, confidence scores were attributed as follows:

**A 1) or A 2) + B(*i*) or B(*ii*) or C(*iii*) = 4**

**A 1) +** **A 2) = 5**

**A 1) + A 2) + B(*i*) or B(*ii*) or C(*iii*)= 7**

| Statistic | Formula used to obtain *r* |
| --- | --- |
| ***t*** | $\sqrt{\frac{t^{2}}{t^{2}+df}}$ |
| ***F*** | $\sqrt{\frac{{df}_{n}F}{{df}_{n}F+{df}_{d}}}$ |
| ***χ2*** | $\sqrt{\frac{\chi^{2}}{N}}$ |
| ***Hedges’ g*** | $\sqrt{\frac{g^{2}n_{1}n_{2}}{g^{2}n_{1}n_{2}+\left( n_{1}n_{2} \right)df}}$ |
| ***R²*** | $\sqrt{\frac{1-(\left( n-1 \right)(1-R^{2}))}{n-k-1}}$ |

**Table S2.** Formulae used to convert statistical values found in initial publications into *r* correlation coefficient (see Nakagawa and Cuthill 2007; Koricheva et al. 2013).

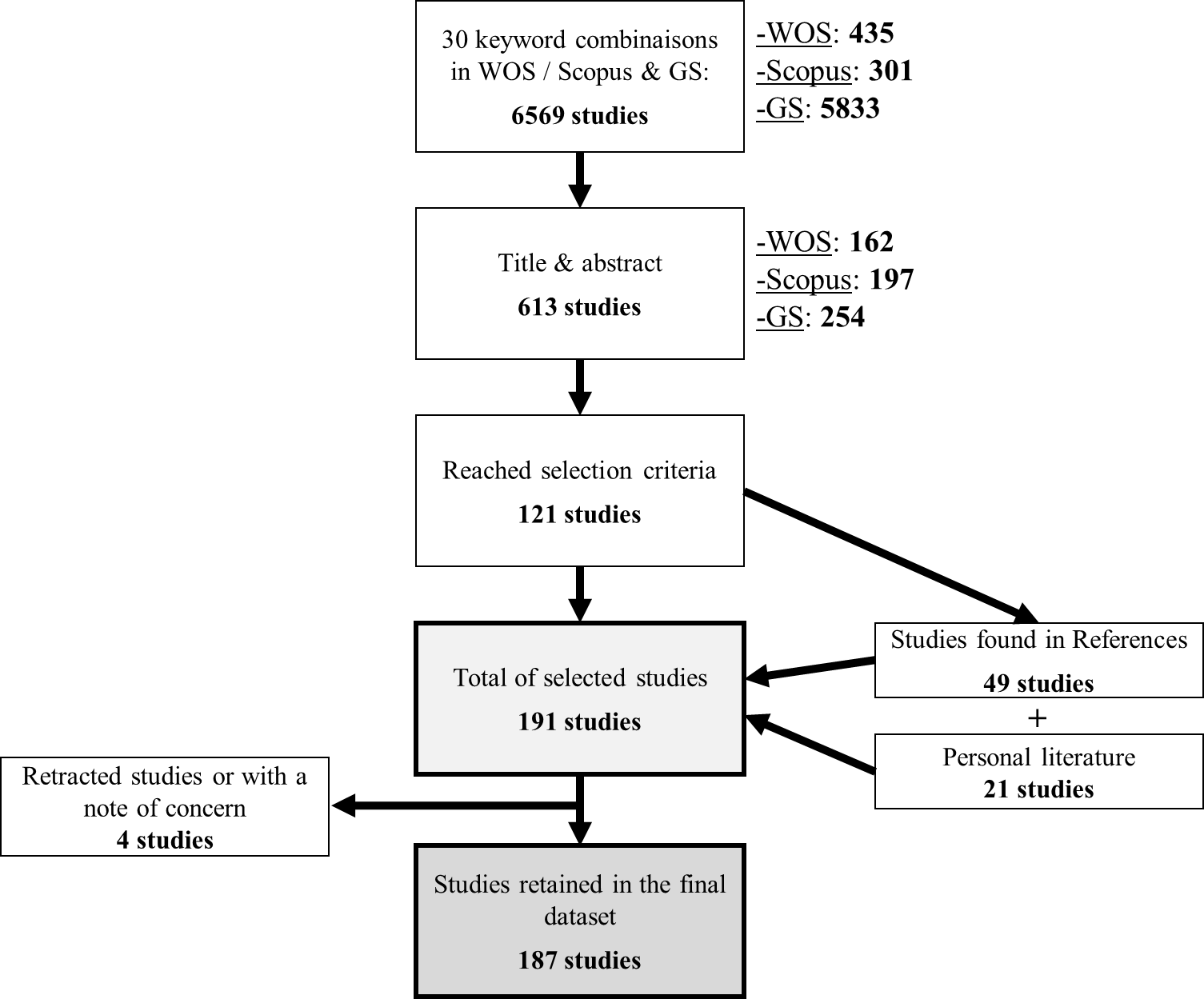

**Figure S1**: Path of the workflow of the systematic review. We used three search engines (WOS: Web Of Science; Scopus; GS: Google Scholar) for each of 30 combinations of keywords (Table S1). We considered the first 200 results from GS for each search term combination. We stopped the systematic review on 29th May 2020. Results at the two first stage of the literature search are results without the duplicates.

**Results associated with the analysis of the whole dataset**

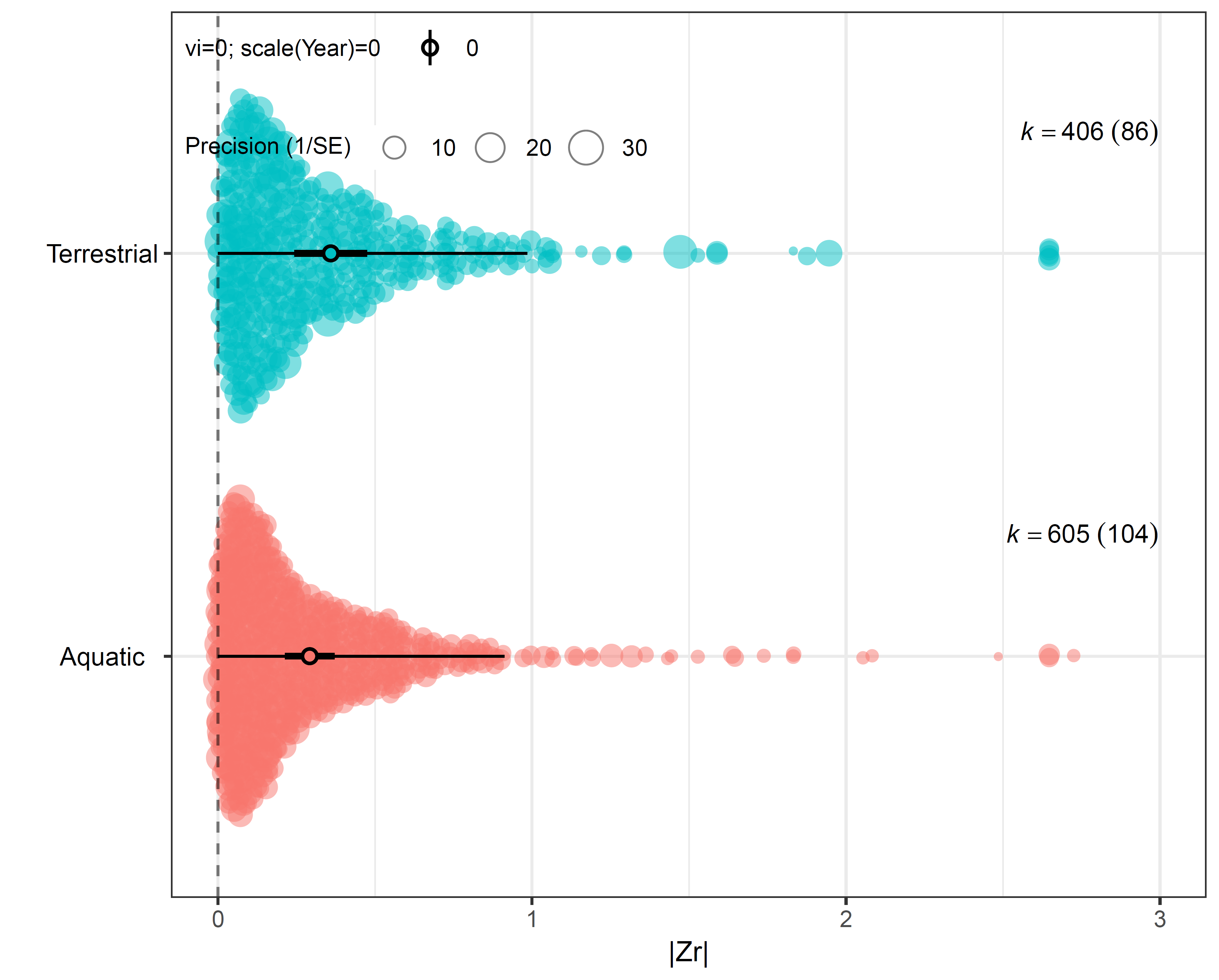

**Figure S2**: Orchard plot of raw effect sizes and mean estimates, 95% confidence intervals (bold error bars), and 95% prediction intervals (error bars) for aquatic and terrestrial realms. The size of bubbles is proportional to their precision (1/SE).

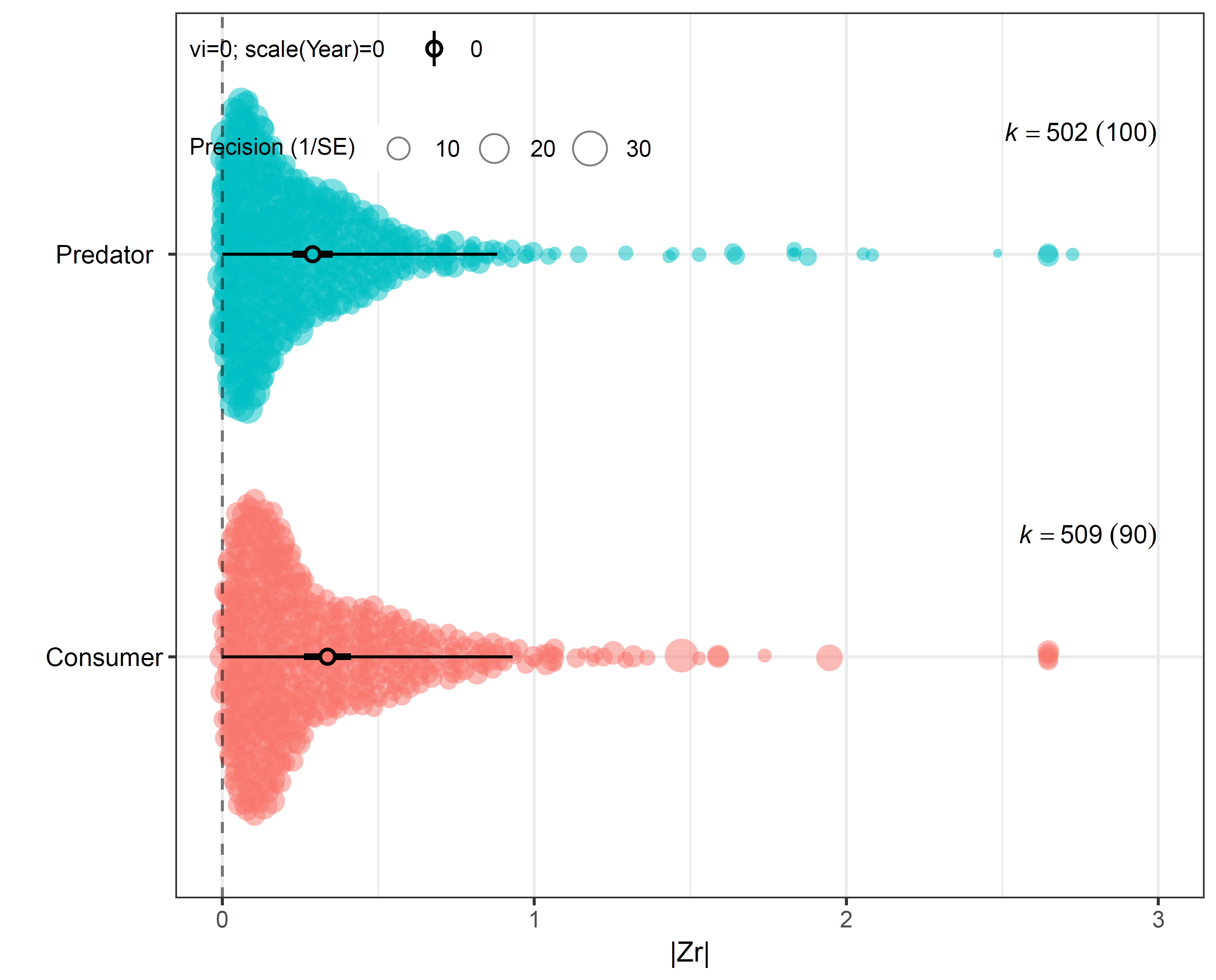

**Figure S3**: Orchard plot of raw effect sizes and mean estimates, 95% confidence intervals (bold error bars), and 95% prediction intervals (error bars) for predators and consumers. The size of bubbles is proportional to their precision (1/SE).

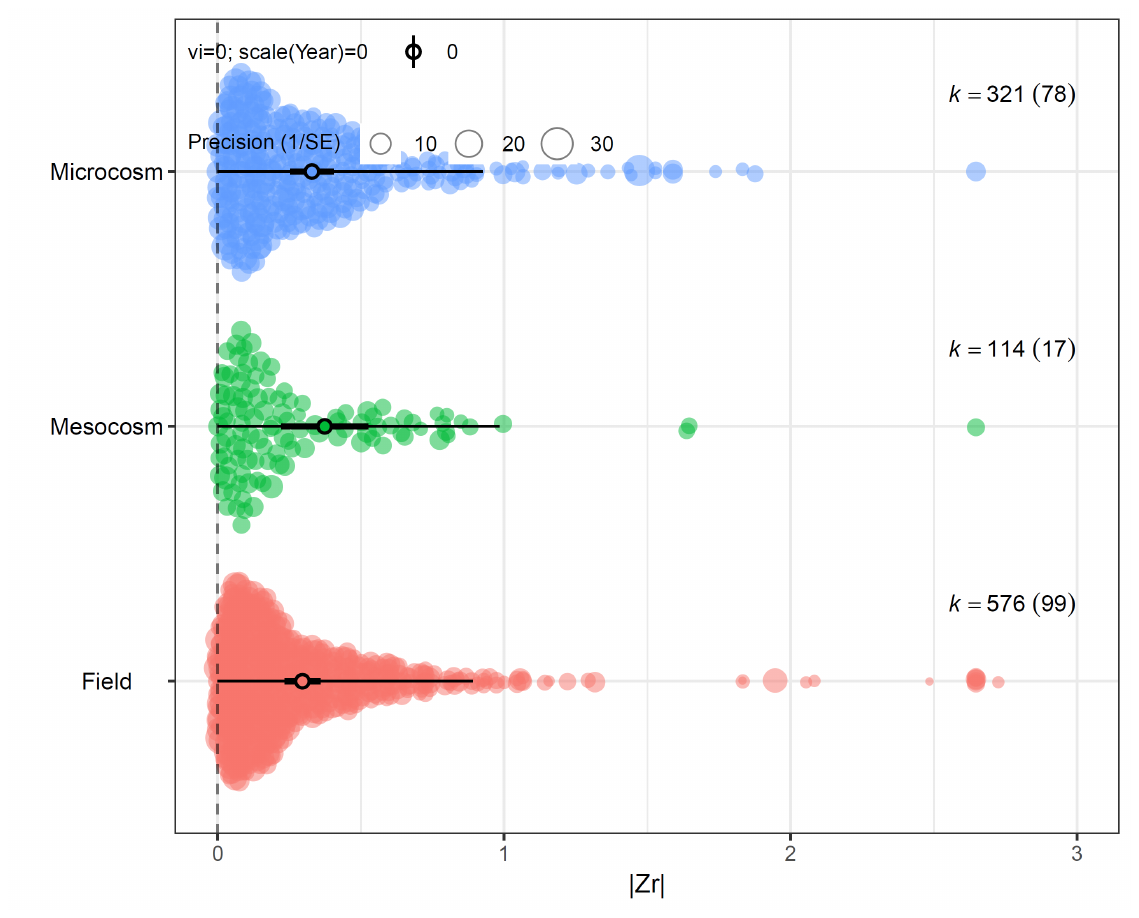

**Figure S4**: Orchard plot of raw effect sizes and mean estimates, 95% confidence intervals (bold error bars), and 95% prediction intervals (error bars) for microcosm, mesocosm, and field settings. The size of bubbles is proportional to their precision (1/SE).

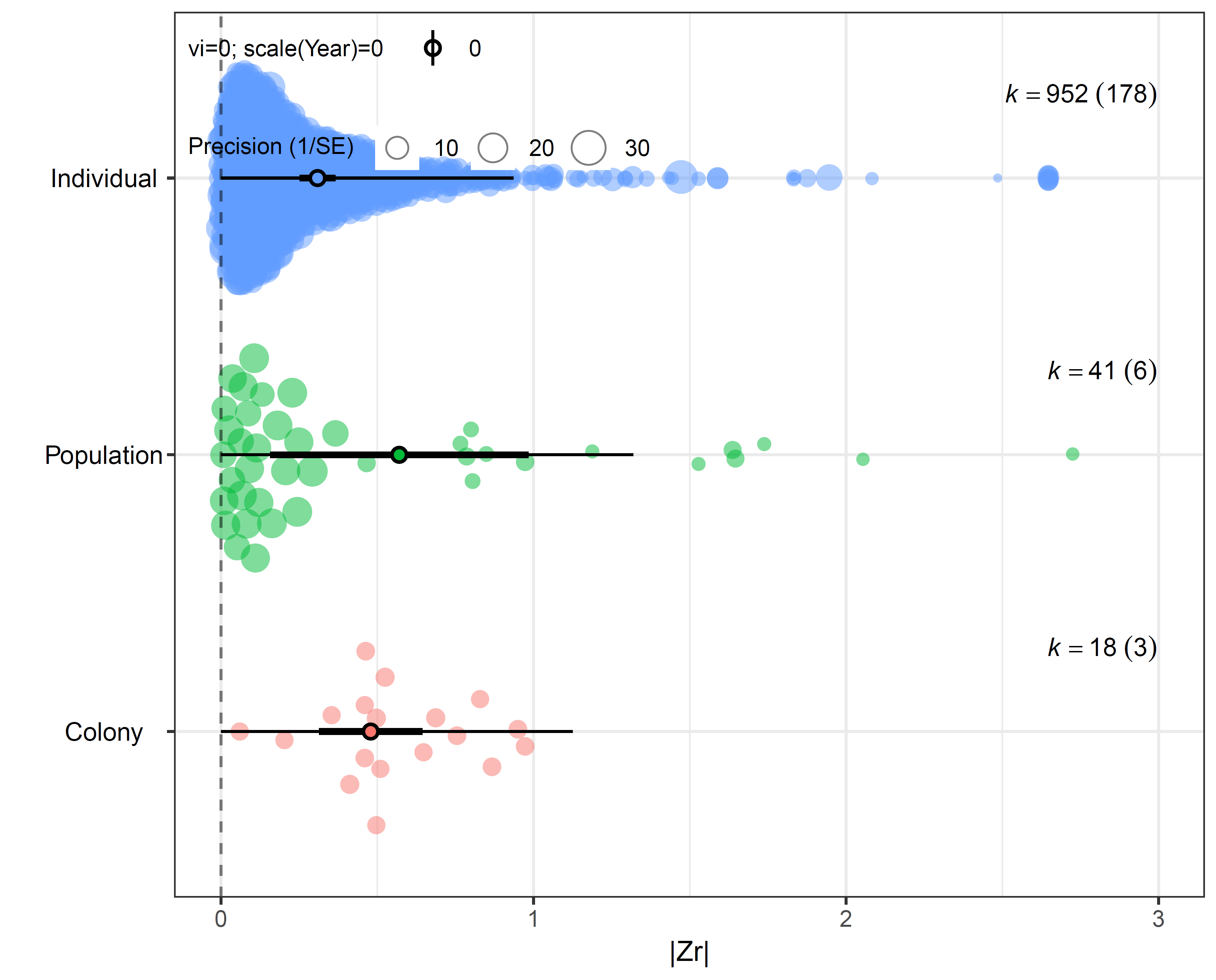

**Figure S5**: Orchard plot of raw effect sizes and mean estimates, 95% confidence intervals (bold error bars), and 95% prediction intervals (error bars) for colonies of eusocial insects, populations, and individuals within populations. The size of bubbles is proportional to their precision (1/SE).

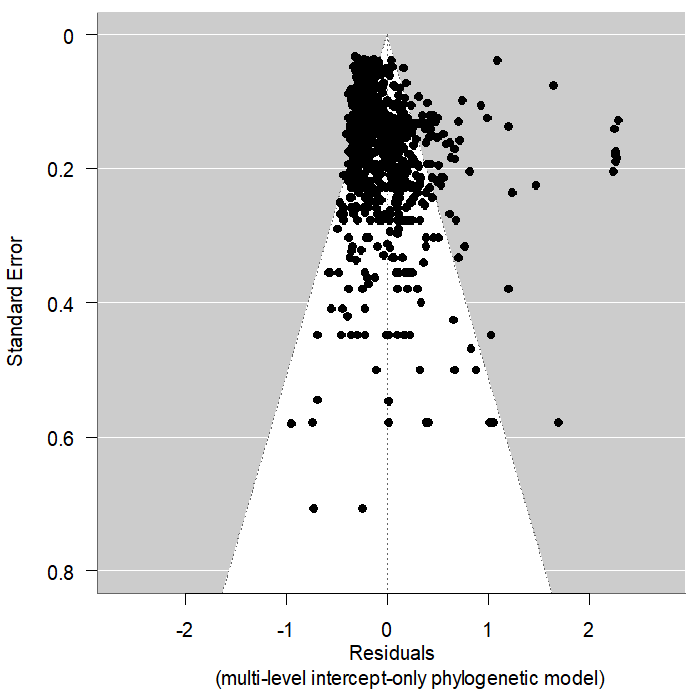

**Figure S6**: Funnel plot of the precision (1/SE) and the residuals of effect sizes |Z*r*| from the intercept-only, phylogenetically-corrected multi-level model. White area shows the pseudo-confidence area at 95%. The grey shaded area indicates where publication bias may occur.

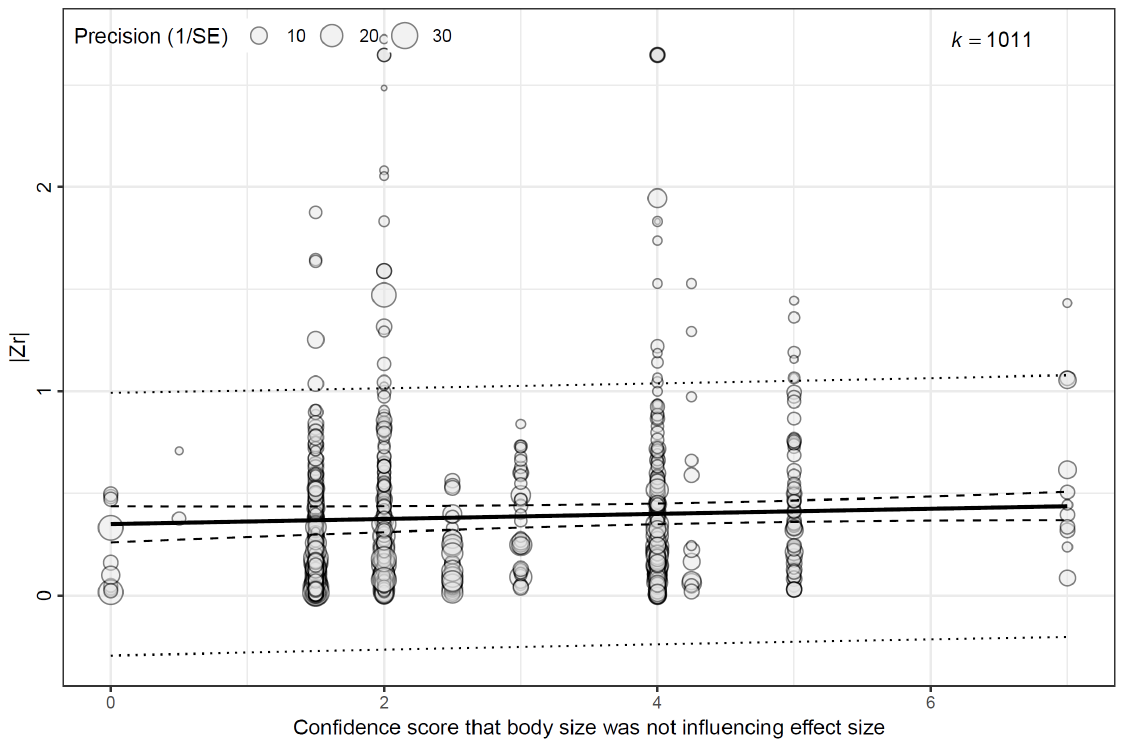

**Figure S7**: Bubble meta-regression plot of the non-significant relationship between effect sizes (|Zr|) and the confidence score (from zero to seven) that we gave for each study regarding the independence (high score values), or the potential for bias (low values) between our effect sizes and with body size (see Box S1, Section 1). Regression line is in solid black, 95% confidence intervals are shown as dashed lines, and prediction intervals as dotted lines. Detail of each score and key element are given in the dataset associated with the paper.

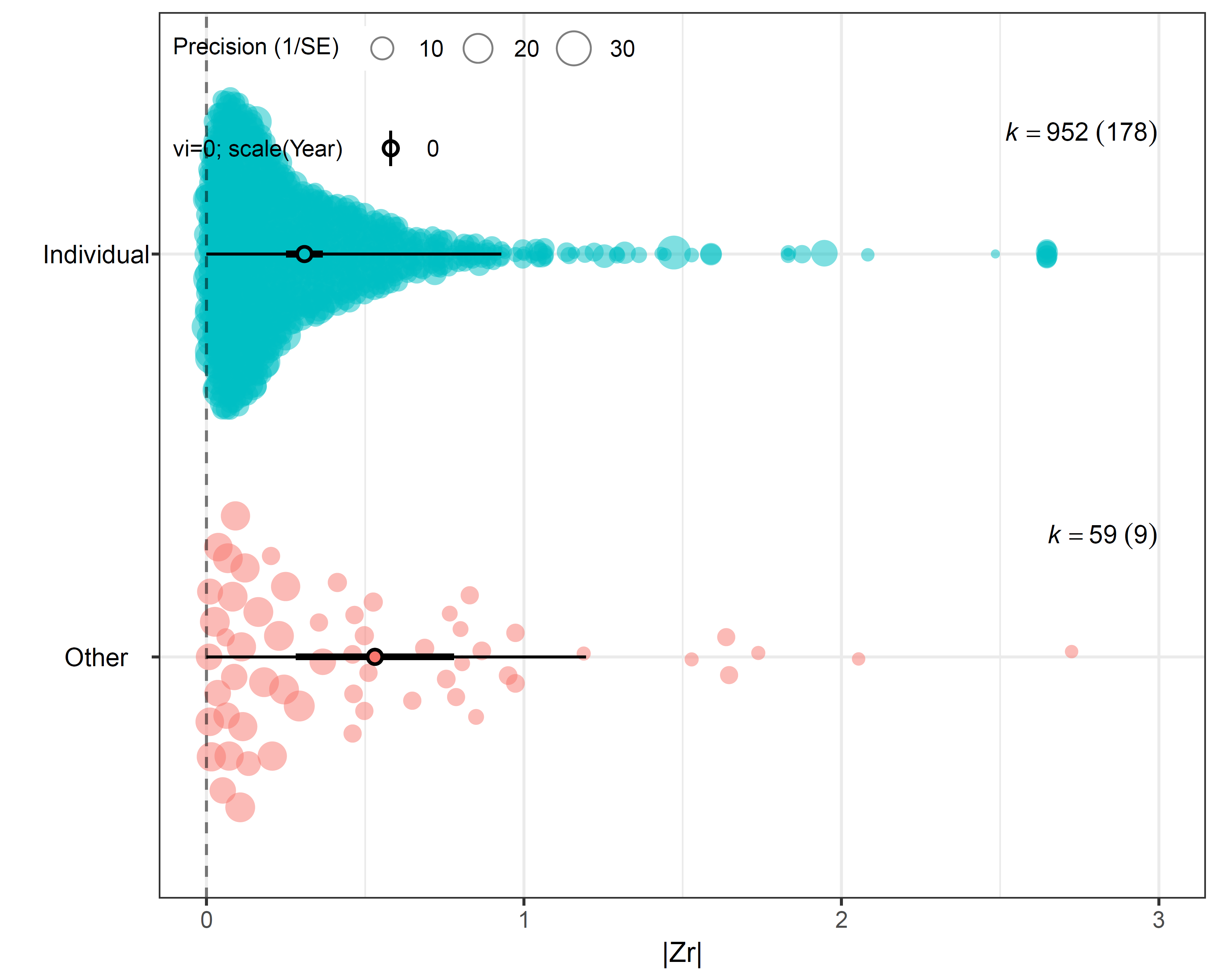

**Figure S8**: Orchard plot of model estimates of |*Zr*| effects sizes estimated for each intraspecific level, i.e., individual (in blue), vs. population and colonies (noted “Other”, in pink), regarding *Hypothesis 1*. The size of each point is proportional to the precision of the effect size (1/SE). Thick and thin error bars give 95% confidence and prediction intervals, respectively. Sample sizes (k) and number of studies (in brackets) are given for each category of ecological responses. Model estimates are reported in table 2, in the main text.

**Table S3**: Table of statistics for the whole dataset (including effect sizes at the individual and at the population levels). Mean effect sizes and 95% confidence intervals are given for response type and trait type models. Raw estimated effect sizes are given as |*Zr*| as well as with the unbiased estimates (|*r*| unbiased). Confidence intervals at 95% are given in brackets. For each model, (**†**) indicates the category with the lowest estimated effect size, and the categories shown in bold (*****) are those with significantly higher estimates compared to (**†**). Pairwise comparison statistics (*z*- and *P*- values) are given accordingly, in comparison to the category with the lowest estimated effect size (**†**). Tests are one-tailed for trait type comparisons (*H3*), two-tailed for response type comparisons (*H2*).

|  | Parameter *(/model)* | \|*Zr*\| | \|*r*\|_unbiased_ | z | *P* |
| --- | --- | --- | --- | --- | --- |
|  | *(/Trait types)* |  |  |  |  |
|  | Morphology **(†)** | 0.29 (0.23 – 0.34) | 0.23 (0.18 – 0.28) | – | – |
|  | Physiology | 0.33 (0.26 – 0.40) | 0.27 (0.21 – 0.32) | 1.26 | .1030 |
|  | **Behavior** **(*)** | **0.33 (0.27 – 0.39)** | **0.27 (0.22 – 0.31)** | **1.85** | **.0319** |
|  | *(/Response types)* |  |  |  |  |
|  | Foraging | 0.34 (0.24 – 0.45) | 0.28 (0.20 – 0.36) | 1.90 | .0570 |
|  | **Trophic niche (*)** | **0.35 (0.25 – 0.45)** | **0.29 (0.21 – 0.37)** | **2.06** | **.0393** |
|  | Growth | 0.34 (0.25 – 0.44) | 0.28 (0.20 – 0.35) | 1.89 | .0594 |
|  | Survival | 0.29 (0.21 – 0.37) | 0.24 (0.17 – 0.30) | 1.34 | .1798 |
|  | Reproduction **(†)** | 0.23 (0.16 – 0.30) | 0.18 (0.12 – 0.24) | – | – |
|  | **Community (*)** | **0.43 (0.28 – 0.59)** | **0.34 (0.21 – 0.46)** | **2.45** | **.0143** |
|  | Ecosystem | 0.41 (0.24 – 0.57) | 0.31 (0.17 – 0.45) | 1.94 | .0519 |

**Table S4:** Checklist of preferred reported items for systematic reviews and meta-analysis in ecology and evolution (PRISMA Eco-evo).

| **Checklist item** | **Sub-item number** | **Description** | **Reported** | **Comment** | **Section** |
| --- | --- | --- | --- | --- | --- |
| **Title and abstract** | 1.1 | Identify the review as a systematic review, meta-analysis, or both | Yes | (…) A meta-analysis | Title  Front page |
|  | 1.2 | Summarise the aims and scope of the review | Yes |  | Abstract |
|  | 1.3 | Describe the data set | Yes |  | Abstract, Introduction, Results |
|  | 1.4 | State the results of the primary outcome | Yes |  | Abstract |
|  | 1.5 | State conclusions | Yes |  | Conclusion |
|  | 1.6 | State limitations | Yes |  | Discussion |
| **Aims and questions** | 2.1 | Provide a rationale for the review | Yes |  | Introduction |
|  | 2.2 | Reference any previous reviews or meta-analyses on the topic | Yes | We referenced to previous syntheses on the topic aiming at describing the magnitude of ITV in plants and animals | Introduction |
|  | 2.3 | State the aims and scope of the review (including its generality) | Yes |  | Introduction |
|  | 2.4 | State the primary questions the review addresses (e.g. which moderators were tested) | Yes |  | Introduction |
|  | 2.5 | Describe whether effect sizes were derived from experimental and/or observational comparisons | Yes | We tested three categories (observational, mesocosm and microcosm) | Methods |
| **Review registration** | 3.1 | Register review aims, hypotheses (if applicable), and methods in a time-stamped and publicly accessible archive and provide a link to the registration in the methods section of the manuscript. Ideally registration occurs before the search, but it can be done at any stage before data analysis. | No | We did not registered our hypotheses before the analysis. | Introduction and methods |
|  | 3.2 | Describe deviations from the registered aims and methods | – |  |  |
|  | 3.3 | Justify deviations from the registered aims and methods | – |  |  |
| **Eligibility criteria** | 4.1 | Report the specific criteria used for including or excluding studies when screening titles and/or abstracts, and full texts, according to the aims of the systematic review (e.g. study design, taxa, data availability) | Yes | PRISMA diagram included as Section 1, Figure S1 and selection criteria in Methods | Methods |
|  | 4.2 | Justify criteria, if necessary (i.e. not obvious from aims and scope) | Yes |  | Methods |
| **Finding studies** | 5.1 | Define the type of search (e.g. comprehensive search, representative sample) | Yes | Representative sample, with 3 different search engines | Methods |
|  | 5.2 | State what sources of information were sought (e.g. published and unpublished studies, personal communications) | Yes | Data came from studies published in ecology and evolution journals | Methods, References |
|  | 5.3 | Include, for each database searched, the exact search strings used, with keyword combinations and Boolean operators | Yes | See Section 1, Table S1 and main text | Methods & Sup. Mat. |
|  | 5.4 | Provide enough information to repeat the equivalent search (if possible), including the timespan covered (start and end dates) | Yes | We provide information on the search engines and keyword settings | Methods & Sup. Mat. |
| **Study selection** | 6.1 | Describe how studies were selected for inclusion at each stage of the screening process (e.g. use of decision trees, screening software) | Yes | See the Figure S1 | Sup. Mat. |
|  | 6.2 | Report the number of people involved and how they contributed (e.g. independent parallel screening) | Yes | See Authorship | Authorship |
| **Data collection process** | 7.1 | Describe where in the reports data were collected from (e.g. text or figures) | Yes | Text | Methods |
|  | 7.2 | Describe how data were collected (e.g. software used to digitize figures, external data sources) | Yes | From the text | Methods |
|  | 7.3 | Describe moderator variables that were constructed from collected data (e.g. number of generations calculated from years and average generation time) | Not applicable |  |  |
|  | 7.4 | Report how missing or ambiguous information was dealt with during data collection (e.g. authors of original studies were contacted for missing descriptive statistics, and/or effect sizes were calculated from test statistics) | Not applicable | We did not include studies for which data were incomplete |  |
|  | 7.5 | Report who collected data | Yes |  | Authorship |
|  | 7.6 | State the number of extractions that were checked for accuracy by co-authors | No |  |  |
| **Data items** | 8.1 | Describe the key data sought from each study | No | We described only a few studies (see Table 1). | Introduction |
|  | 8.2 | Describe items that do not appear in the main results, or which could not be extracted due to insufficient information | Not applicable |  |  |
|  | 8.3 | Describe main assumptions or simplifications that were made (e.g. categorizing both 'length' and 'mass' as 'morphology') | Yes |  | Methods |
|  | 8.4 | Describe the type of replication unit (e.g. individuals, broods, study sites) | Yes |  | Methods |
| **Assessment of individual study quality** | 9.1 | Describe whether the quality of studies included in the systematic review or meta-analysis was assessed (e.g. blinded data collection, reporting quality, experimental versus observational) | Yes | We included a covariate for the study design (observational, mesocosm, microcosm) | Methods |
|  | 9.2 | Describe how information about study quality was incorporated into analyses (e.g. meta-regression and/or sensitivity analysis) | Yes | We acknowledged and controlled for point estimates bias in adding sampling variances (*vi*) as covariates. | Methods, Statistical analysis section |
| **Effect size measures** | 10.1 | Describe effect size(s) used | Yes |  | Methods |
|  | 10.2 | Provide a reference to the equation of each calculated effect size (e.g. standardized mean difference, log response ratio) and (if applicable) its sampling variance | Yes | We referred to Nagakawa et al. (2007), that we followed to calculate Z*r* values from different statistics. Sampling variances (*vi*) were calculated with the R package ‘*metafor*’. | Methods |
|  | 10.3 | If no reference exists, derive the equations for each effect size and state the assumed sampling distribution(s) | Not applicable |  |  |
| **Missing data** | 11.1 | Describe any steps taken to deal with missing data during analysis (e.g. imputation, complete case, subset analysis) | Not applicable |  |  |
|  | 11.2 | Justify the decisions made to deal with missing data | Not applicable |  |  |
| **Meta-analytic model description** | 12.1 | Describe the models used for synthesis of effect sizes | Yes | Hierarchical multi-level phylogenetic meta-analytic models (function ‘*rma.mv*’ in ‘*metafor*’ package in R) | Methods |
|  | 12.2 | The most common approach in ecology and evolution will be a random-effects model, often with a hierarchical/multilevel structure. If other types of models are chosen (e.g. common/fixed effects model, unweighted model), provide justification for this choice | Not applicable | We used a hierarchical multi-level model (see above) |  |
| **Software** | 13.1 | Describe the statistical platform used for inference (e.g. R) | Yes | R | Methods |
|  | 13.2 | Describe the packages used to run models | Yes | (‘metafor’ in R) | Methods |
|  | 13.3 | Describe the functions used to run models | Yes | We used ‘*rma.mv’* |  |
|  | 13.4 | Describe any arguments that differed from the default settings | Yes | Codes and data are available | Methods  Sup. Mat. |
|  | 13.5 | Describe the version numbers of all software used | Yes |  | Methods |
| **Non-independence** | 14.1 | Describe the types of non-independence encountered (e.g. phylogenetic, spatial, multiple measurements over time) | Yes |  | Methods |
|  | 14.2 | Describe how non-independence has been handled | Yes |  | Methods |
|  | 14.3 | Justify decisions made | Yes |  | Methods |
| **Meta-regression and model selection** | 15.1 | Provide a rationale for the inclusion of moderators (covariates) that were evaluated in meta-regression models | Yes |  | Methods |
|  | 15.2 | Justify the number of parameters estimated in models, in relation to the number of effect sizes and studies (e.g. interaction terms were not included due to insufficient sample sizes) | Yes |  | Methods |
|  | 15.3 | Describe any process of model selection | Not applicable | We only ran models that were of interest for our hypotheses. So we did not perform model selection | Methods |
| **Publication bias and sensitivity analysis** | 16.1 | Describe assessments of the risk of bias due to missing results (e.g. publication, time-lag, and taxonomic biases) | Yes | See according paragraphs regarding the statistical methods and the report of the results (publication bias assessment) | Methods, Results |
|  | 16.2 | Describe any steps taken to investigate the effects of such biases (if present) | Yes | We implemented several steps to investigate the effects of bias. These were random effects accounting for the multi-level non-independence of effect sizes (within and among studyID), a robust estimation of effect sizes with a variance-covariance matrix acknowledging for sources of non-independences in the dataset, acknowledging for effect size precision (sampling variance as a fixed effect), species phylogeny, a null model, time-lag with year of publication as a fixed effect (please see the statistical analysis sub-section in the Methods section) | Methods and Results |
|  | 16.3 | Describe any other analyses of robustness of the results, e.g. due to effect size choice, weighting or analytical model assumptions, inclusion or exclusion of subsets of the data, or the inclusion of alternative moderator variables in meta-regressions | Yes | We unbiased effect sizes in computing null effect sizes (effect sizes expected under the null hypothesis of no effect)  We fit the results on two datasets. One including the full dataset (H1), and one including only observations among individuals (H2 and H3). Results for H2 and H3 on the full dataset are given in Sup. Mat., which led to qualitatively similar conclusions | Methods, Results, and Discussion |
| **Clarification of post hoc analyses** | 17.1 | When hypotheses were formulated after data analysis, this should be acknowledged. | Yes | We designed hypotheses before the data analyses. Additional analyses that were designed a posteriori are labelled as post hoc analyses (i.e., Figure 5). | Methods, statistical analysis section and results |
| **Metadata, data, and code** | 18.1 | Share metadata (i.e. data descriptions) | Yes | See link to data | Data accessibility |
|  | 18.2 | Share data required to reproduce the results presented in the manuscript | Yes | We share the data used to perform the statistics reported in the paper | See link to data |
|  | 18.3 | Share additional data, including information that was not presented in the manuscript (e.g. raw data used to calculate effect sizes, descriptions of where data were located in papers) | Yes |  | See link to data |
|  | 18.4 | Share analysis scripts (or, if a software package with graphical user interface (GUI) was used, then describe full model specification and fully specify choices) | Yes | We share the R code used to perform the statistics reported in the paper | See link to data |
| **Results of study selection process** | 19.1 | Report the number of studies screened | Yes | see Figure S1 | Sup. Mat. |
|  | 19.2 | Report the number of studies excluded at each stage of screening | Yes | see Figure S1 | Sup. Mat. |
|  | 19.3 | Report brief reasons for exclusion from the full text stage | Yes | The study does not fill our selection criteria | Methods |
|  | 19.4 | Present a Preferred Reporting Items for Systematic Reviews and Meta-Analyses (PRISMA)-like flowchart (www.prisma-statement.org). | Yes | see Figure S1 | Sup. Mat. |
| **Sample sizes and study characteristics** | 20.1 | Report the number of studies and effect sizes for data included in meta-analyses | Yes |  | Results |
|  | 20.2 | Report the number of studies and effect sizes for subsets of data included in meta-regressions | Yes | see Orchard plot figures and Tables in main text | Results |
|  | 20.3 | Provide a summary of key characteristics for reported outcomes (either in text or figures; e.g. one quarter of effect sizes reported for vertebrates and the rest invertebrates) | Yes | see Orchard plot figures and Tables in main text | Results |
|  | 20.4 | Provide a summary of limitations of included moderators (e.g. collinearity and overlap between moderators) | Yes |  | Discussion |
|  | 20.5 | Provide a summary of characteristics related to individual study quality (risk of bias) | No |  |  |
| **Meta-analysis** | 21.1 | Provide a quantitative synthesis of results across studies, including estimates for the mean effect size, with confidence/credible intervals | Yes | Main text | Results |
| **Heterogeneity** | 22.1 | Report indicators of heterogeneity in the estimated effect (e.g. I2, tau2 and other variance components) | Yes | I² at all hierarchical levels, Q-statistic on total heterogeneity, see methods on statistics and results in main text regarding heterogeneity assessment | Results |
| **Meta-regression** | 23.1 | Provide estimates of meta-regression slopes (i.e. regression coefficients) and confidence/credible intervals | Yes | Table 2, Figure 3–5, Main text | Results |
|  | 23.2 | Include estimates and confidence/credible intervals for all moderator variables that were assessed (i.e. complete reporting) | Yes | Table 2, Figure 3–5, Main text | Results, Sup. Mat. |
|  | 23.3 | Report interactions, if they were included | Not applicable | No interactions were evaluated |  |
|  | 23.4 | Describe outcomes from model selection, if done (e.g. R2 and AIC) | Not applicable | We did not perform model selection |  |
| **Outcomes of publication bias and sensitivity analysis** | 24.1 | Provide results for the assessments of the risks of bias (e.g. Egger's regression, funnel plots) | Yes | Main text and figures in Sup. Mat. | Methods, Results and Sup. Mat. |
|  | 24.2 | Provide results for the robustness of the review's results (e.g. subgroup analyses, meta-regression of study quality, results from alternative methods of analysis, and temporal trends) | Yes | We analysed main hypotheses on different subsets of data, with very little variation overall, indicating robust results | see Methods, Results, Sup. Mat. |
| **Discussion** | 25.1 | Summarise the main findings in terms of the magnitude of effect | Yes | Table 2, Figure 3–5, Main text | Results, Discussion |
|  | 25.2 | Summarise the main findings in terms of the precision of effects (e.g. size of confidence intervals, statistical significance) | Yes | Table 2, Figure 3–5, Main text | Results, Discussion |
|  | 25.3 | Summarise the main findings in terms of their heterogeneity | Yes | I², main text | Results |
|  | 25.4 | Summarise the main findings in terms of their biological/practical relevance | Yes |  | Results, Discussion and Conclusion |
|  | 25.5 | Compare results with previous reviews on the topic, if available | Yes |  | Discussion |
|  | 25.6 | Consider limitations and their influence on the generality of conclusions, such as gaps in the available evidence (e.g. taxonomic and geographical research biases) | Yes |  | Discussion |
| **Contributions** | 26.1 | Provide names, affiliations, and funding sources of all co-authors | Yes |  | Front page |
|  | 26.2 | List the contributions of each co-author | Yes |  | Authorship |
|  | 26.3 | Provide contact details for the corresponding author | Yes |  | Front page |
|  | 26.4 | Disclose any conflicts of interest | Yes | We have no conflict of interest to declare | Conflict of interest statement |
| **References** | 27.1 | Provide a reference list of all studies included in the systematic review or meta-analysis | Yes |  | References for Meta-Analysis |
|  | 27.2 | List included studies as referenced sources (e.g. rather than listing them in a table or supplement) | Yes |  | References for Meta-Analysis |

**References cited in Section 1**

Koricheva, J., J. Gurevitch, and K. Mengersen, eds. 2013. Handbook of meta-analysis in ecology and evolution. Princeton University Press, Princeton.

Moiron, M., K. L. Laskowski, and P. T. Niemelä. 2020. Individual differences in behaviour explain variation in survival: a meta‐analysis. (J. Gurevitch, ed.) Ecology Letters 23:399–408.

Nakagawa, S., and I. C. Cuthill. 2007. Effect size, confidence interval and statistical significance: a practical guide for biologists. Biological Reviews 82:591–605.

Smith, B. R., and D. T. Blumstein. 2008. Fitness consequences of personality: a meta-analysis. Behavioral Ecology 19:448–455.

***Section 2****: Ad hoc analysis of the strength of relationships between individual variability in body size and various ecological responses*

We aimed to give an element of comparison to our estimates of the strength of relationships between traits varying the most independently from body size or ontogeny, in giving a meta-analytical estimate of the strength of relationships belonging to individual differences in body size. Rather than performing a systematic search here (which would require a dedicated study), we looked for published studies or meta-analyses on that specific issue.

***Foraging –*** We identified four studies reporting estimates of relationships between size and ecological responses covering the spectrum of ecological responses that we synthetized in our main analysis. First, we extracted correlation coefficients from Maino and Kearney (2015), and from Rota et al. (2022). From these two studies, we extracted 48 effect sizes of relationships between body mass (dry weight) and feeding rates estimated at the individual level, from approximately a similar number of insect and arthropod species.

***Fitness –*** Assessing the links between ontogenetic variation in body size (among different age or size classes) and fitness does not seem to have been an important aspect of evolutionary ecology studies. It is trivial that mortality is elevated for older individuals, and that immature individuals have no reproduction outputs. Perhaps as a result, we had difficulty to find relevant and consequent datasets for this issue. However, in Ronget et al. (2018), the authors performed a thorough meta-analysis on the effect of offspring body mass variation (that can vary substantially as they nicely introduce in their paper), on survival, advocating that selection pressure on juvenile survival is a key element of the life-history of animal populations (they focused on birds and mammals). We collected all their 161 effect sizes (i.e., odd ratios that we converted to Cohen’s *d*, and then to *Zr* effect sizes), from more than a hundred of species.

***Community & ecosystem –*** We found relevant the design of Rudolf and Rasmussen (2013), who tested in small ponds how the body size or ontogeny of aquatic predators (a beetle and a dragonfly) affected the community of aquatic prey and lower trophic levels, and key ecosystem processes (respiration, NPP and decomposition). From this study, we collected the 18 effect sizes presented in Table 1 of their paper (i.e., “stage” and “stage-by-species” Chi² statistics on each dependent response). One treatment in this study was mixing small and large groups of individuals, but as the effects were mainly additive, the global effect sizes from this study remain relevant to our question (while potentially leading to an overestimation of effect sizes).

***Description and analysis of the ad hoc dataset –*** This leads us to an *ad hoc* dataset of 227 effect sizes of the relationship strengths of individual variability in body mass with various ecological responses, i.e., feeding rate, fitness (survival), and community/ecosystem related responses. The data belonged from 163 studies, and 139 species of animals (among birds, mammals, insects, and arthropods). As the great majority of relationships were naturally positive (more than 95% of effect sizes), we did not apply our null model correction approach here. Proportions of sample sizes for each of these categories (21%, 71% and 8%, for foraging, fitness, and community/ecosystem, respectively) were similar to the ones of our main dataset (26%, 57% and 17%, for foraging, fitness, and community/ecosystem, respectively). We treated these ad hoc data separately, but in a similar way than our main dataset. We started by fitting a similar multi-level intercept-only model, including “*vi”* and study year (centered to zero) as fixed covariates. We simplified the model structure, and replaced the phylogenetic approach by random effects of ‘*Taxonomic group*’ (either mammals, birds, arthropods, or insects) nested in ‘*Species*’. Similarly, to account for variability among source datasets, studies, and effect sizes within studies, we nested ‘*within-studyID*’ within ‘*StudyID*’ within ‘*meta-datasetID*’. We also added a random effect for the type of ‘*ecological responseID*’ in our intercept-only model that we used to estimate the grand-mean effect size. The overall heterogeneity for the intercept-only model was moderate, with a total heterogeneity of 58.5%, and the funnel plot of the residuals of this model appeared overall symmetric, suggesting only few instances of publication bias (Figure S9). We then removed the latter random effect in the second model that we used to test differences among ecological responses (the same term was passed as a fixed effect).

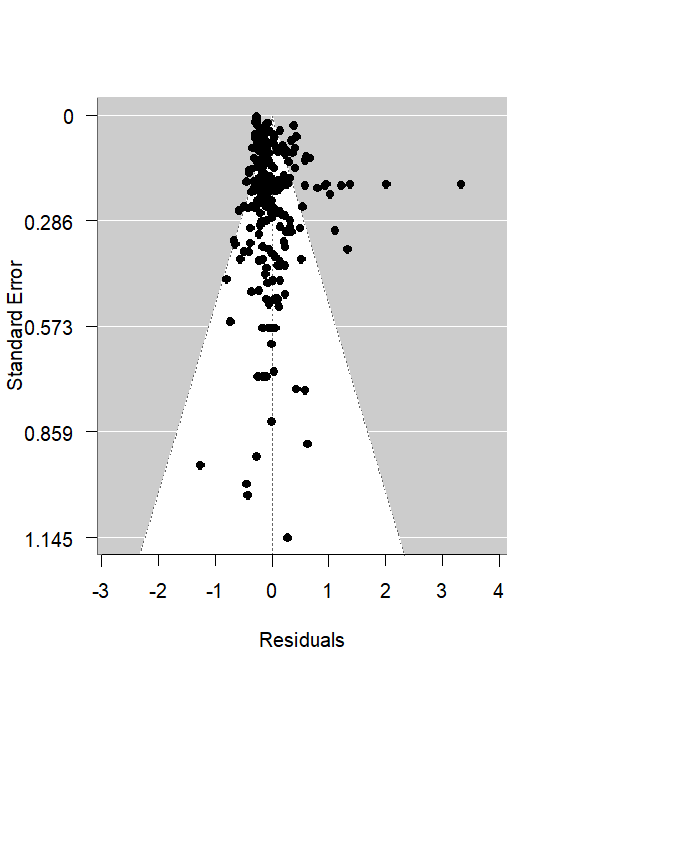

**Figure S9**. Funnel plot of the residuals of the ad hoc multi-level model. White area depicts the pseudo-confidence area at 95% where effect sizes suffer small publication bias, the grey area shows effect sizes potentially suffering from publication bias.

***Results –*** Surprisingly, and contrary to our expectation, we found that overall, effect sizes of relationships of body size varying among individuals (|*r*| = 0.28 [0.19 – 0.37] 95% CIs), were largely comparable to the ones we previously estimated from other individual-level phenotypic traits assessed on similarly sized individuals, or for which potential body size effects were accounted for (|*r*|_unbiased_ = 0.26 [0.21 – 0.30] 95% CIs; given in main text).

Interestingly, on this independent ad hoc dataset, we found that the strength of relationships maintained by individual variation in body size we estimated for each of the three categories of ecological responses was following qualitatively our results regarding phenotypic traits varying beyond body size; with higher effect sizes for foraging (|*r*| = 0.36 [0.22 – 0.48] 95% CIs) and community/ecosystem responses (|*r*| = 0.73 [0.71 – 0.76] 95% CIs), than for survival (|*r*| = 0.25 [0.17 – 0.32] 95% CIs; Figure S10). Effect sizes at community/ecosystem levels were significantly higher than those for foraging and survival (*z* = 22.95; *P* < .0001 and *z* = 10.52; *P* < .0001, respectively), and effect sizes for foraging were also higher than those estimated for survival (*z* = 1.99; *P* = .0463).

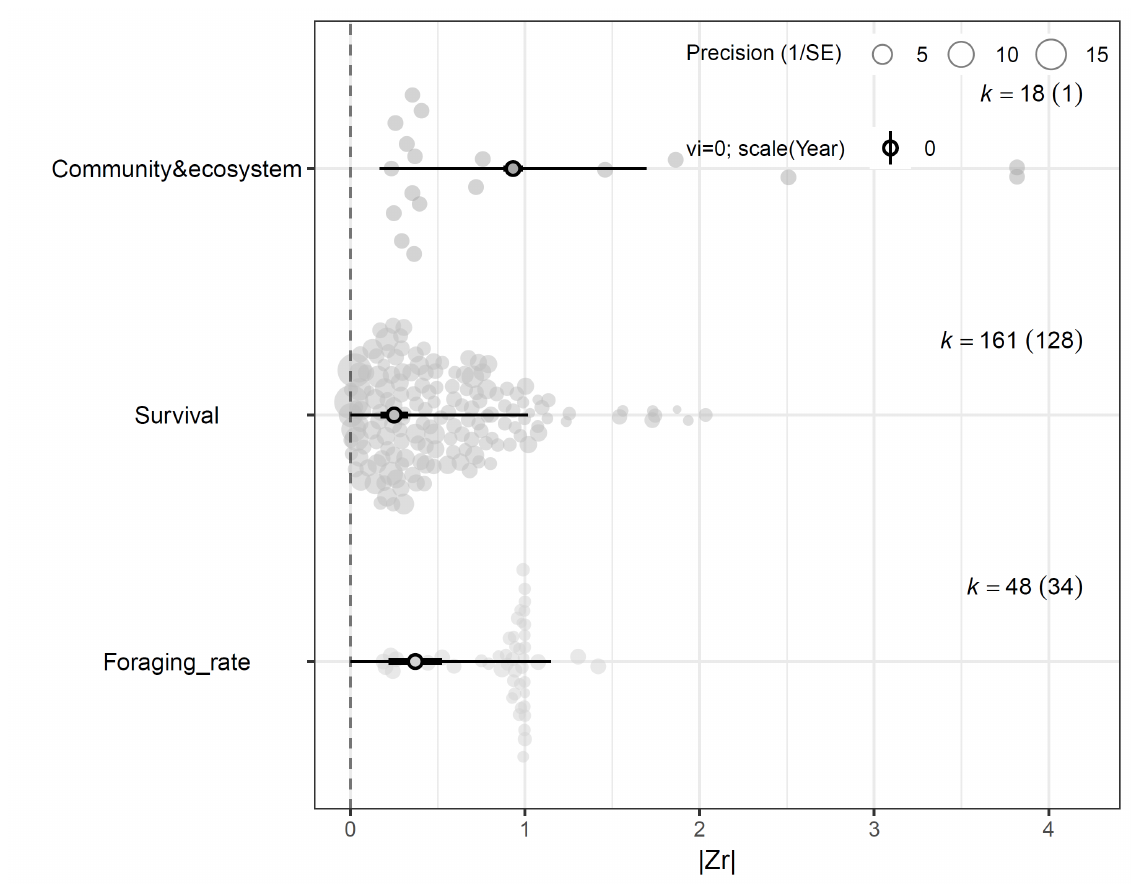

**Figure S10**. Orchard plot showing |*Zr*| effect sizes, point estimates, confidence intervals at 95% and prediction intervals at 95% (solid and thin bars, respectively) for each ecological response category of the ad hoc dataset of body size-to-ecological correlations at the individual level.

**References cited in Section 2**

Maino, J. L., and M. R. Kearney. 2015. Ontogenetic and interspecific scaling of consumption in insects. Oikos 124:1564–1570.

Ronget, V., J. Gaillard, T. Coulson, M. Garratt, F. Gueyffier, J. Lega, and J. Lemaître. 2018. Causes and consequences of variation in offspring body mass: meta‐analyses in birds and mammals. Biological Reviews 93:1–27.

Rota, T., A. Lecerf, É. Chauvet, and B. Pey. 2022. The importance of intraspecific variation in litter consumption rate of aquatic and terrestrial macro-detritivores. Basic and Applied Ecology 63:175–185.

Rudolf, V. H. W., and N. L. Rasmussen. 2013. Population structure determines functional differences among species and ecosystem processes. Nature Communications 4:2318.
